# Rapid Emergence and Evolution of SARS-CoV-2 Variants in Advanced HIV Infection

**DOI:** 10.1101/2024.01.05.574420

**Authors:** Sung Hee Ko, Pierce Radecki, Frida Belinky, Jinal N. Bhiman, Susan Meiring, Jackie Kleynhans, Daniel Amoako, Vanessa Guerra Canedo, Margaret Lucas, Dikeledi Kekana, Neil Martinson, Limakatso Lebina, Josie Everatt, Stefano Tempia, Tatsiana Bylund, Reda Rawi, Peter D. Kwong, Nicole Wolter, Anne von Gottberg, Cheryl Cohen, Eli A. Boritz

## Abstract

Previous studies have linked the evolution of severe acute respiratory syndrome coronavirus-2 (SARS-CoV-2) genetic variants to persistent infections in people with immunocompromising conditions^1–4^, but the evolutionary processes underlying these observations are incompletely understood. Here we used high-throughput, single-genome amplification and sequencing (HT-SGS) to obtain up to ∼10^3^ SARS-CoV-2 spike gene sequences in each of 184 respiratory samples from 22 people with HIV (PWH) and 25 people without HIV (PWOH). Twelve of 22 PWH had advanced HIV infection, defined by peripheral blood CD4 T cell counts (i.e., CD4 counts) <200 cells/μL. In PWOH and PWH with CD4 counts ≥200 cells/μL, most single-genome spike sequences in each person matched one haplotype that predominated throughout the infection. By contrast, people with advanced HIV showed elevated intra-host spike diversity with a median of 46 haplotypes per person (IQR 14-114). Higher intra-host spike diversity immediately after COVID-19 symptom onset predicted longer SARS-CoV-2 RNA shedding among PWH, and intra-host spike diversity at this timepoint was significantly higher in people with advanced HIV than in PWOH. Composition of spike sequence populations in people with advanced HIV fluctuated rapidly over time, with founder sequences often replaced by groups of new haplotypes. These population-level changes were associated with a high total burden of intra-host mutations and positive selection at functionally important residues. In several cases, delayed emergence of detectable serum binding to spike was associated with positive selection for presumptive antibody-escape mutations. Taken together, our findings show remarkable intra-host genetic diversity of SARS-CoV-2 in advanced HIV infection and suggest that adaptive intra-host SARS-CoV-2 evolution in this setting may contribute to the emergence of new variants of concern (VOCs).

## Main

While mounting evidence suggests that SARS-CoV-2 genetic variants emerge preferentially in immunocompromised individuals, the processes of intra-host evolution that produce these variants are incompletely understood. Multiple studies have documented new SARS-CoV-2 mutations in people with HIV (PWH), with conditions requiring immunosuppressive therapy, and/or with B cell deficiencies^5–12^. New mutations have typically been detected in these cases weeks or months after COVID-19 symptom onset, with one recent study^13^ reporting that overall SARS-CoV-2 mutation rates were similar between short-term and persistent infections. These findings suggest a temporal threshold after which SARS-CoV-2 has accumulated enough mutations to evolve within the individual. However, convergent evolution of the same mutations in unrelated persistent cases implies extensive early intra-host SARS-CoV-2 sequence diversification that has not been directly observed. Many persistent infections described in previous studies have been characterized retrospectively, with limited analysis during the acute phase. Equally important, while standard technologies can track SARS-CoV-2 consensus sequence changes and identify some minor variant mutations in genomic surveillance^14–17^, advanced approaches that define intra-host virus genetic diversity and evolution at the single-genome level have not been widely used. Addressing these gaps may help elucidate how SARS-CoV-2 establishes persistent infection and generates new sequence variants.

To define the genetic diversity and evolutionary signatures among SARS-CoV-2 genomes in each individual, we have developed a high-throughput, single-genome amplification and sequencing (HT-SGS) approach that combines unique barcoding of virus genomes with long-read sequencing to produce up to ∼10^3^ single-copy sequences per sample^18^. Here we used HT-SGS of the full-length spike gene to analyze a unique cohort of clinically diverse PWH and people without HIV (PWOH) sampled from onset to clearance of SARS-CoV-2 infection^19,20^. Through longitudinal analysis of SARS-CoV-2 spike sequences together with detection of anti-spike antibody binding, we find unique aspects of SARS-CoV-2 evolution in people with advanced, poorly controlled HIV infection that markedly increase the risk for generation of new SARS-CoV-2 variants in these individuals.

### Longitudinal sampling of PWH and PWOH

We investigated intra-host evolution of SARS-CoV-2 during persistent infection using longitudinal sample sets from 22 PWH and 25 PWOH. These individuals had participated in cohort studies of people with COVID-19 diagnosed either as hospital inpatients or as outpatients between May 1, 2020 and December 31, 2020 (hospitalized cohort)^19^ or between October 2, 2020 and September 30, 2021 (outpatient cohort)^20^. From the hospitalized cohort, we included a subgroup of 10 PWH with peripheral blood CD4 T cell counts (i.e., CD4 counts) <200 cells/μL who had high initial SARS-CoV-2 RNA levels in respiratory samples (rRT-PCR cycle threshold [Ct] <30) (**Extended Data Fig. 1; Supplementary Table 1)**. Remaining participants from the hospitalized cohort included 5 PWH for whom CD4 counts were not available, 3 PWH with CD4 counts ≥200 cells/μL, and 7 PWOH. From the outpatient cohort we included 2 PWH with CD4 counts <200 cells/μL, 2 PWH with CD4 counts ≥200 cells/μL, and 18 PWOH **(Extended Data Fig. 1a; Supplementary Table 1)**. Notably, among the 12 PWH with CD4 counts <200 cells/μL, 6 had plasma HIV RNA >10^5^ copies/mL, and 4 had no plasma HIV RNA level documented **(Supplementary Table 1)**. Upper respiratory tract samples were available from these individuals beginning at study enrollment, at a median of 4 days (IQR 3-8) after the onset of COVID-19 symptoms, and every second day (hospitalized cohort) or three times weekly (outpatient cohort) thereafter until the cessation of SARS-CoV-2 RNA shedding. As described previously in the hospitalized cohort^19^, PWH with CD4 counts <200 cells/μL who had high initial SARS-CoV-2 RNA levels in respiratory specimens often experienced prolonged SARS-CoV-2 RNA shedding **(Extended Data Fig. 1b)**.

### HT-SGS vs. standard whole-genome sequencing

SARS-CoV-2 spike sequences in upper respiratory tract samples from these individuals were determined by HT-SGS of the full-length spike gene. This approach detected mutations that were present in as few as 0.45% of amplifiable virus genomes per sample **(Extended Data Fig. 2a)**. HT-SGS demonstrates mutational linkage patterns across the 3.8-kilobase spike region that are not detectable by short-read whole-genome sequencing (WGS) **(Extended Data Fig. 2b-d)**. By defining and quantifying the unique linked groupings of mutations (i.e., haplotypes) in each sample at the level of single-genome sequences (SGS), HT-SGS provides minimum estimates of intra-host population diversity and enables downstream analysis of evolutionary relationships among viruses in each person **(Extended Data Fig. 2b-d)**.

### Intra-host SARS-CoV-2 genetic diversity in PWH and PWOH

We used HT-SGS to analyze 184 samples from the 47 study participants, resulting in 70,968 SGS from PWH and 29,824 SGS from PWOH. These SGS included 431 different single-nucleotide variations (SNVs) or deletions that together defined 831 spike gene haplotypes. As shown in **Fig. 1a**, strikingly high numbers of spike haplotypes were detected in some PWH. This was especially true for PWH with CD4 counts <200 cells/μL (sky blue tree sections, **Fig. 1a**), in whom a median of 46 haplotypes/person (IQR 14-114/person) were detected over the course of infection. Analysis of very rare SNVs and deletions indicated the presence of additional haplotypes at levels below reportable limits at the given sampling depth, particularly in PWH with CD4 counts <200 cells/μL **(Extended Data Fig. 3a)**. Considering each sample timepoint separately to avoid inflating genetic-distance-based diversity calculations in prolonged infections, we found higher intra-host diversity in PWH with CD4 counts <200 cells/μL than in PWH with higher CD4 counts or in PWOH as measured by normalized Shannon entropy, average pairwise genetic distance, and total numbers of haplotypes detected **(Fig. 1b)**. These diversity measures were similar between PWH with higher CD4 counts and PWOH. While the number of haplotypes identified in each sample was positively correlated with the number of SGS obtained from that sample **(Extended Data Fig. 3b)**, ratios of haplotypes identified per SGS obtained were nonetheless significantly higher in PWH with CD4 counts <200 cells/μL than in the other subgroups **(Extended Data Fig. 3c)**. SARS-CoV-2 RNA levels were largely independent of diversity measures within subgroups **(Extended Data Fig. 3d)**. Moreover, the differences in SARS-CoV-2 spike gene diversity between subgroups described above were also observed in an analysis limited to the hospitalized cohort **(Extended Data Fig. 3e)**. We conclude that HT-SGS revealed a markedly elevated intra-host diversity of SARS-CoV-2 spike haplotypes among people with advanced HIV infection.

**Fig. 1.**
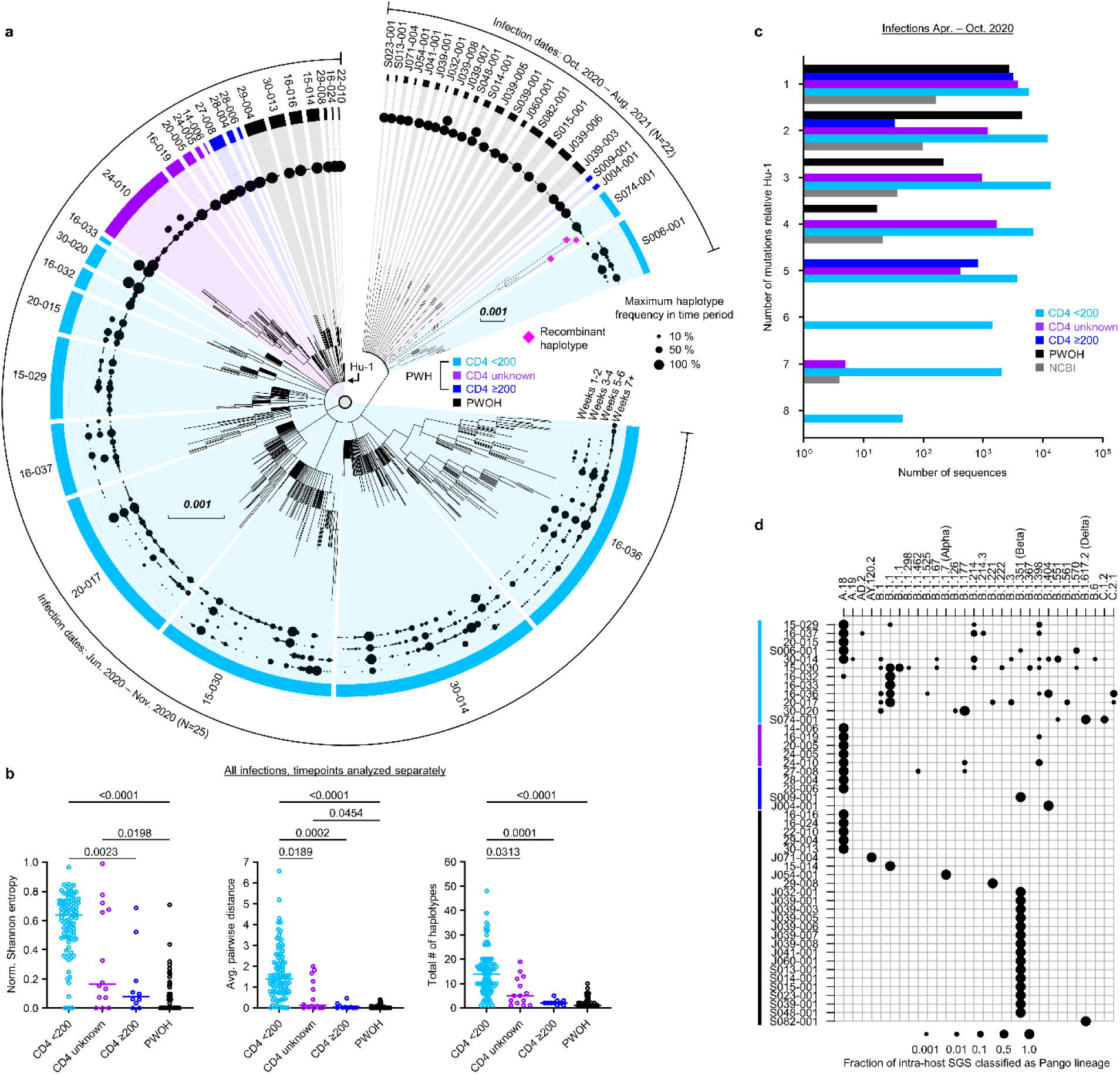
Intra-host diversity of SARS-CoV-2 spike sequences in PWH and PWOH. (a) Maximum-likelihood phylogenetic analysis of spike gene haplotypes detected in each participant. Trees were generated separately for each participant and then joined for visualization. The maximum frequency of each haplotype during each two-week period is shown with a scaled black dot. Trees from hospitalized and outpatient cohorts are separated to reflect differences in infecting Pango lineages. Color coding of PWOH and PWH subgroups applies to all figures. (b) Comparison of spike genetic diversity among PWOH and subgroups of PWH. Individual samples from longitudinal sample sets in each person are represented by separate datapoints. Statistical significance was assessed by one-way ANOVA with multiple comparisons (Kruskal-Wallis test and Dunn’s multiple comparisons test); *p* values <0.05 are shown. (c) Genetic divergence (number of mutations, y-axis) from ancestral Wuhan-Hu-1 in SARS-CoV-2 spike sequences from PWH subgroups, PWOH, and matched public data. Data are shown for infections detected between April and October 2020. Numbers of sequences (x-axis) indicate total numbers of single-genome sequences (SGS) for all participants combined in PWOH and PWH subgroups, or total numbers of single-person consensus sequences obtained from the NCBI Virus database (grey). Statistical significance was assessed by one-way ANOVA with multiple comparisons (Friedman test and Dunn’s multiple comparisons test); statistically significant *p* values were detected for the following comparisons: PWH CD4 <200 vs. NCBI (*p* = 0.005), PWH CD4 <200 vs. PWH CD4 ≥200 (*p* = 0.002), and PWH CD4 <200 vs. PWOH (*p* = 0.006). (d) Analysis of secondary intra-host Pango lineages. Distinct lineages are indicated as columns; individual participants are indicated as rows. Dots represent relative frequencies of individual lineages among all SGS from each participant.

To relate SARS-CoV-2 sequence variation detected in our study participants to the variation among viruses circulating contemporaneously in the same geographic region, we compared spike genetic divergence from Wuhan-Hu-1 (Hu-1, GenBank Accession NC_045512.2, nucleotide coordinates 21563-25384) between SGS from the hospitalized cohort and matched public WGS data^21^ **(Fig. 1c)**. In data on the NCBI Virus database from infections in South Africa between April and October 2020, 91.7% of spike sequences showed 1-3 mutations relative to Hu-1. In contrast, 31% of spike SGS from PWH with CD4 counts <200 cells/μL had 4 or more mutations relative to Hu-1 (*p* = 0.005 for comparison of distributions in PWH with CD4 counts <200 cells/μL vs. NCBI, Friedman test with Dunn’s multiple comparisons test). Furthermore, SARS-CoV-2 sequence diversification in PWH from both hospitalized and outpatient cohorts was associated with intra-host emergence of variants that mapped by Nextclade^22^ to secondary Pango lineages **(Fig. 1d)**. Among PWH with CD4 counts <200 cells/μL, this analysis indicated a median of 3.5 lineages (range, 1-9) in each person. Secondary intra-host lineages thus identified in PWH with CD4 counts <200 cells/μL included B.1.1.525, B.1.214.3, and C.2.1 at timepoints preceding the first reports of these lineages in global surveillance (1/5/2021, 12/14/2020, and 12/18/2020). No secondary Pango lineages were identified among intra-host sequences from PWOH, regardless of infecting variant or clinical cohort **(Fig. 1d)**. We conclude that SARS-CoV-2 sequence divergence from ancestral detected by HT-SGS in people with advanced HIV significantly exceeded the divergence of geographically-and temporally matched circulating sequences, in some cases anticipating the later emergence of SARS-CoV-2 genetic variants in the general population.

### Spike evolution over time

To define the kinetics of intra-host SARS-CoV-2 evolution in our study cohort, we analyzed longitudinal patterns in spike HT-SGS data from PWH and PWOH. We observed that normalized Shannon entropy, average pairwise genetic distance, and total haplotype numbers at the first sample timepoint after symptom onset were significantly higher in PWH with CD4 counts <200 cells/μL than in PWOH **(Fig. 2a)** even though days between symptom onset and the first sample timepoint were not significantly different between groups (PWH with CD4 counts <200 cells/μL median 4.5 days, IQR 4-13.5; PWOH median 3 days, IQR 2-5; *p* = 0.1526, Kruskal-Wallis). Higher initial spike gene diversity predicted longer SARS-CoV-2 RNA shedding duration among PWH **(Fig. 2b)** but not among PWOH **(Extended Data Fig. 3f)**. All measures of intra-host spike gene diversity increased significantly over time among PWH with CD4 counts <200 cells/μL, reflecting longer infections in participants with high initial diversity as well as rising diversity in some participants **(Fig. 2c).** In a longitudinal analysis of the most abundant spike haplotypes in each person, rapid fluctuations in frequency observed in most PWH with CD4 counts <200 cells/μL **(Extended Data Fig. 4a)** contrasted sharply with the persistence of a single predominant haplotype throughout the course of infection in every PWOH **(Extended Data Fig. 4b)**. Haplotype abundance fluctuations were often associated with progressively lower frequencies of the presumptive founder haplotype over time in each PWH with CD4 count <200 cells/μL **(Fig. 2d)**. The use of Jensen-Shannon distance to quantify global changes in population haplotype composition over time in all pairs of sample timepoints in each person revealed large changes in PWH with CD4 counts <200 cells/μL **(Fig. 2e)**. Although the magnitude of these changes was correlated with the time between samples, large changes were observed in PWH with CD4 counts <200 cells/μL even over short time intervals (main panel and inset panel, **Fig. 2e**). Taken together, these findings show that the development of elevated SARS-CoV-2 spike gene diversity in PWH with CD4 counts <200 cells/μL encompasses 1) high early diversity, beginning shortly after COVID-19 symptom onset, and 2) marked and rapid changes in the population of sequences detected in each person over time.

**Fig. 2.**
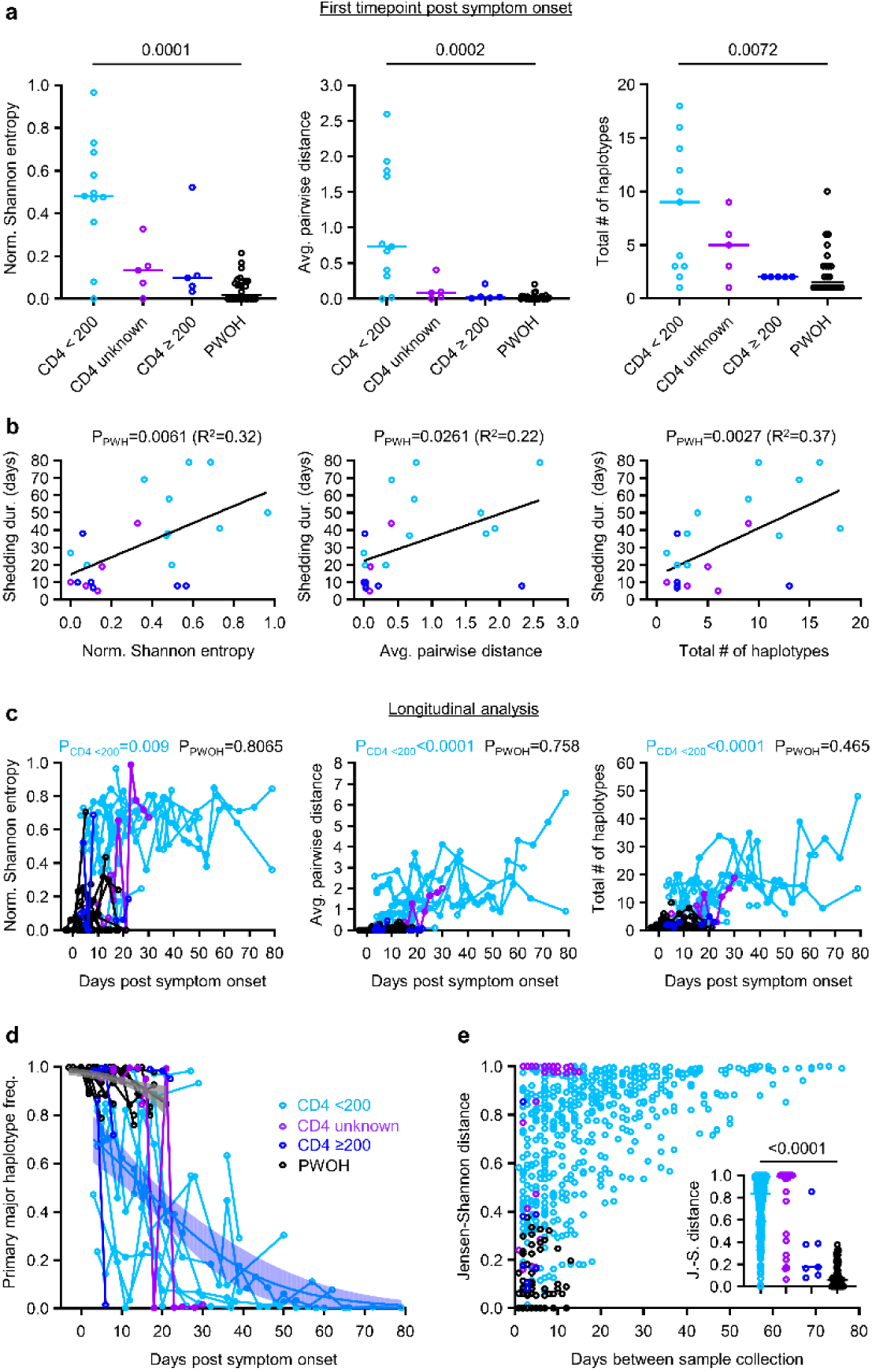
Longitudinal analysis of intra-host spike evolution in PWH and PWOH. (a) Comparison of spike genetic diversity among PWOH and subgroups of PWH at the first sample timepoint. Statistical significance was assessed by one-way ANOVA with multiple comparisons (Kruskal-Wallis test and Dunn’s multiple comparisons test); *p* values <0.05 are shown. (b) Correlations between measurements of spike diversity at the first sample timepoint and SARS-CoV-2 RNA shedding duration in all PWH analyzed together. Linear regression lines are shown. (c) Longitudinal changes in measurements of spike diversity in PWH subgroups and PWOH. Spearman *p* values for correlations between measurements of spike diversity and time of sampling (days post symptom onset) are shown for PWH with CD4 counts <200 cells/μL and PWOH. (d) Longitudinal changes in the intra-host frequency in PWH subgroups and PWOH of the primary major haplotype (i.e., the most abundant haplotype detected in the individual’s first sample timepoint). Blue and grey curves indicate logistic regressions of frequency declines for PWH with CD4 counts <200 cells/μL and PWOH; shaded areas indicate bootstrapped 95% confidence intervals. (e) Pairwise similarity analysis (Jensen-Shannon distance) of virus populations for all pairs of sample timepoints in each participant. The inset panel compares Jensen-Shannon distances by participant subgroup for sample pairs collected ≤14 days apart. Statistical significance was assessed by one-way ANOVA with multiple comparisons (Kruskal-Wallis test and Dunn’s multiple comparisons test).

### Analysis of transmitted SARS-CoV-2 spike diversity

The diversity of spike sequences detected in some PWH at early timepoints after COVID-19 symptom onset raised the possibility of multiple founder sequences in these individuals. To address this possibility, we performed single-linkage phylogenetic clustering to identify clades of haplotypes in each person that were separated by at least 5 mutations and were thus likely to have originated from distinct founders^23^. This analysis revealed evidence of multiple founder sequences in 2 PWH, both of whom had CD4 counts <200 cells/μL **(Extended Data Fig. 5a)**. No evidence for >1 founder sequence was detected in any PWOH. In the PWH with evidence for >1 founder sequence, recombination analysis for participant S074-001 revealed low frequencies of 3 distinct intra-host recombinant haplotypes between Delta (B.1.617.2) and C.1.2 variant lineages **(Extended Data Fig. 5a, b)**. Recombinant haplotypes were not detected in the other participants, consistent with an absence of recombinant haplotypes and/or limitations in identifying recombinant haplotypes in the setting of low genetic diversity. We conclude that infections with multiple SARS-CoV-2 founder sequences can be detected in PWH, and this may contribute to the generation of further virus genetic diversity through intra-host recombination.

### Spike mutations in PWH and PWOH

We investigated the nature of SARS-CoV-2 genetic diversity in PWH and PWOH by compiling all spike gene positions at which SGS were polymorphic over the course of each participant’s infection. Intra-host polymorphisms (i.e., mutations found in <100% of SGS in the person) included synonymous SNVs, nonsynonymous SNVs, and deletions **(Fig. 3)**. Synonymous SNVs were scattered across the spike gene at different positions in different participants **(Fig. 3a, b)**, reflecting their expected evolutionary neutrality, and were found at higher levels in PWH with CD4 counts <200 cells/μL than in the other subgroups **(Fig. 3c)**. These findings were consistent with elevated cumulative numbers of replicative cycles in PWH with CD4 counts <200 cells/μL, as suggested by high initial virus RNA levels and prolonged shedding in these individuals^19^. At the same time, HT-SGS revealed extensive intra-host nonsynonymous spike gene variation in the cohort. Many nonsynonymous mutations were deletions or nonsynonymous SNVs at three recurrently deleted sites in the NH_2_-terminal domain (NTD)^24^, with additional nonsynonymous SNVs in the receptor-binding domain (RBD), furin cleavage site, and membrane fusion regions, and with deletions in the fusion peptide **(Fig. 3a, b)**. Although several PWH with higher CD4 counts and PWOH also showed nonsynonymous intra-host mutations **(Fig. 3a, b)**, the number of nonsynonymous intra-host mutations per person was significantly higher in PWH with CD4 counts <200 cells/μL than in the other subgroups **(Fig. 3d)**. The intra-host nonsynonymous mutations detected in PWH with CD4 counts <200 cells/μL overlapped significantly with the defining mutations of variants of concern (VOCs) Omicron BA.1, Alpha, Beta, and Delta variants of concern (44.2% [23 of 52] of codons with nonsynonymous mutations in VOCs also mutated in PWH with CD4 counts <200 cells/μL, vs. 12.1% [155 of 1273] of all codons in spike mutated in PWH with CD4 counts <200 cells/μL; Fisher’s exact test *p* <0.0001) **(Fig. 3e)**. Moreover, structure-based calculations of solvent-accessible surface area (SASA)^25,26^ for each amino acid residue in spike showed that, while synonymous mutations were not biased to residues with high or low SASA, nonsynonymous mutations were more commonly detected in solvent-accessible residues (**Extended Data Fig. 6a, b)**. The strength of this bias was similar in residues mutated in VOCs (**Extended Data Fig. 6a)**. Thus, HT-SGS of the spike gene showed that well-described mutations associated with worldwide SARS-CoV-2 evolution occurred commonly as intra-host mutations in PWH with CD4 counts <200 cells/μL, in the context of a high total burden of mutations in these individuals.

**Fig. 3.**
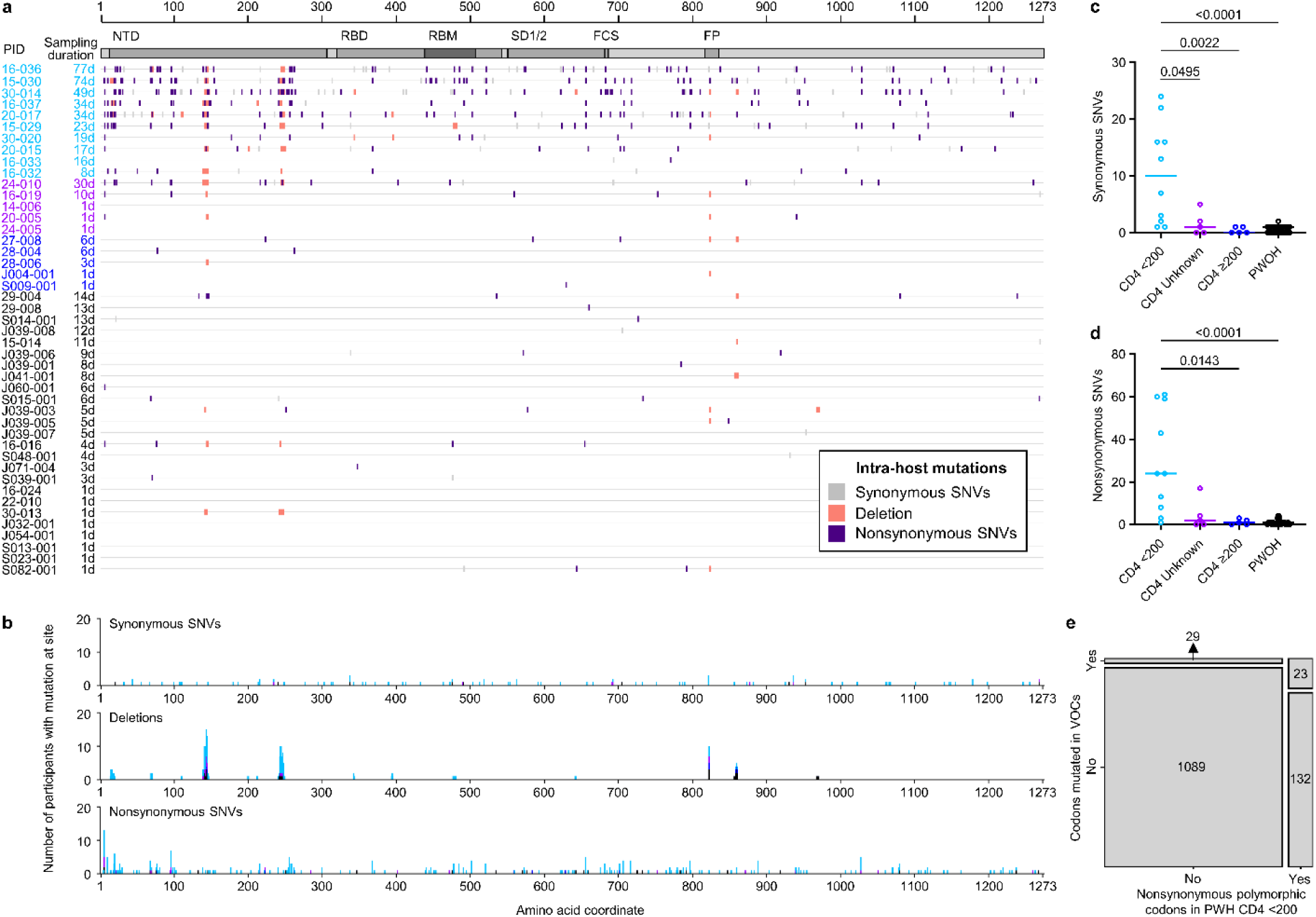
Analysis of intra-host spike mutations in PWH and PWOH. (a) Locations and types of intra-host spike mutations detected over all timepoints in each participant. Participants S006-001 and S074-001 were infected with multiple founders and are not shown. (b) Total numbers of participants with synonymous (top), deletion (middle), and nonsynonymous (bottom) mutations detected at the indicated positions over all timepoints. Stacked bars are colored by participant subgroup. (c and d) Numbers of intra-host synonymous (c) and nonsynonymous (d) single-nucleotide variations (SNVs) by participant, compared among PWH subgroups and PWOH. Statistical significance was assessed by one-way ANOVA with multiple comparisons (Kruskal-Wallis test and Dunn’s multiple comparisons test); *p* values <0.05 are shown. (e) Contingency analysis of codons in spike with nonsynonymous intra-host mutations in PWH with CD4 counts <200 cells/μL vs. codons with nonsynonymous mutations in the VOCs Alpha, Beta, Delta, and/or Omicron BA.1. The association between sites with nonsynonymous mutations in PWH with CD4 counts <200 cells/μL and sites mutated in the VOCs was significant (Fisher’s exact test, *p* <0.00001).

### Selection analysis

We next asked whether patterns of spike gene evolution in PWH with CD4 counts <200 cells/μL indicated adaptation of the virus to the host or were instead consistent with chance expansion of variant haplotypes bearing random mutations (i.e., genetic drift). We used the FUBAR algorithm^27^ in conjunction with the maximum-likelihood phylogeny for each participant to identify codons at which the calculated ratio of nonsynonymous to synonymous mutations (dN/dS) supported positive selective pressure. To minimize false-positive selection analysis results owing to recurrent low-frequency mutations, we considered mutations identified by dN/dS to be under positive selection only if their intra-host frequency increased by >20% during the individual’s infection. This analytical approach demonstrated positive selection at one or more spike gene positions in 7 of the 12 PWH with CD4 counts <200 cells/μL (see colored symbols, **Fig. 4** and **Extended Data Fig. 5a**). Intra-host mutations were identified by selection analysis in these individuals in the signal peptide, NTD, RBD, subdomain 1 and 2, and S2 regions of the spike gene. In sharp contrast, no sites under positive selection were detected in either PWOH **(Extended Data Fig. 7)** or PWH with CD4 counts ≥200 cells/μL **(Extended Data Fig. 8)**. One (1) site under positive selection was detected in a PWH with unknown CD4 count who showed high spike diversity (participant 24-010, see **Extended Data Fig. 8**). Thus, selection analysis enabled by HT-SGS suggested that the intra-host diversity of SARS-CoV-2 spike sequences in PWH with CD4 counts <200 cells/μL arose in part through adaptive evolution, and not purely through genetic drift.

**Fig. 4.**
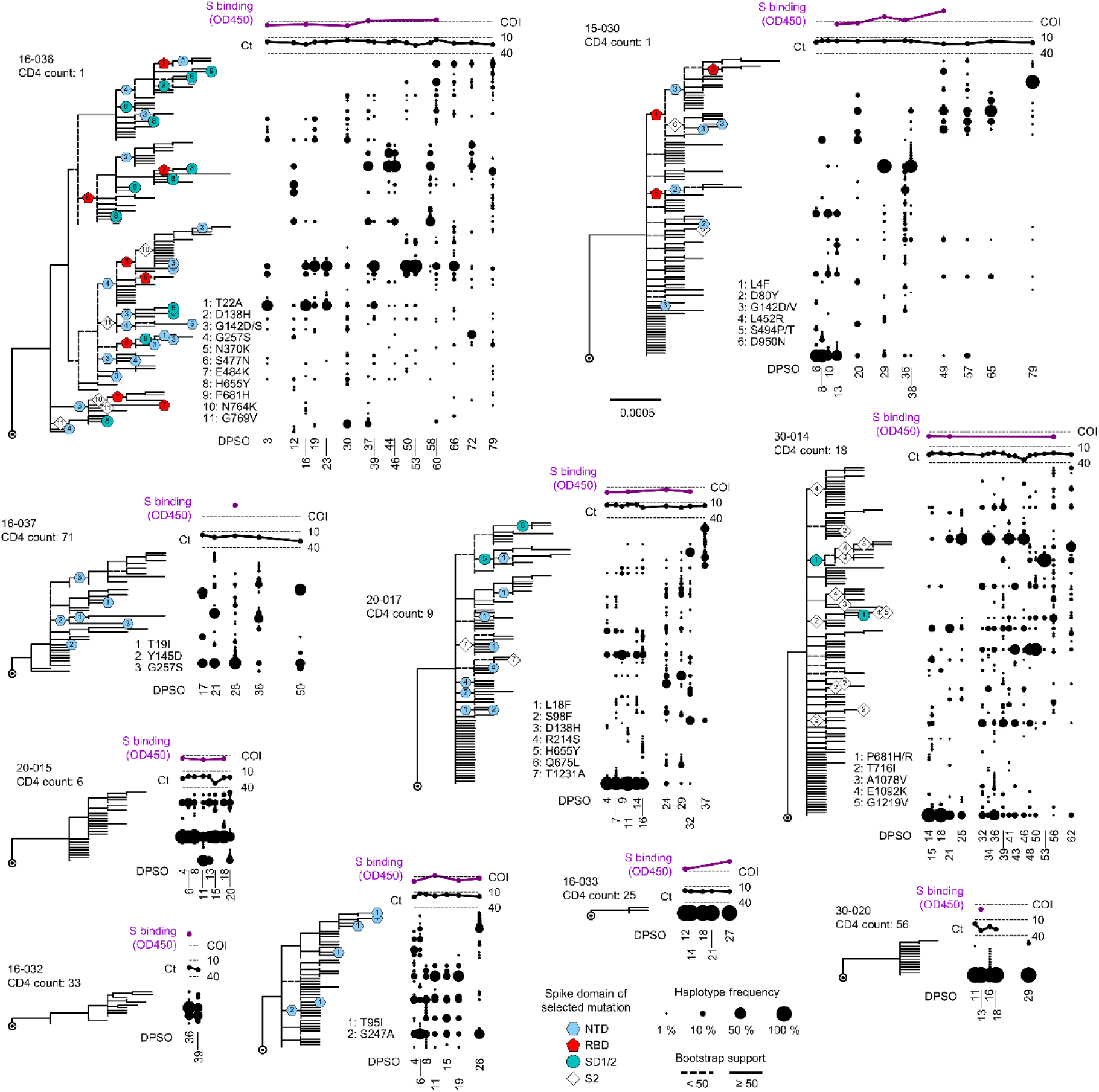
Evolution and positive selection of SARS-CoV-2 spike in PWH with CD4 counts <200 cells/μL. Maximum-likelihood phylogenetic trees rooted on Hu-1 for all haplotypes from each PWH with CD4 counts <200 cells/μL. Participants S006-001 and S074-001 were infected with multiple founders and are showed separately in **Extended Data Fig. 5a.** Clades with bootstrap support <50% are indicated with dashed lines. Sites detected under positive selection within each participant (see Methods) are shown at their inferred location on the tree with numbered symbols; mutations corresponding to each number are listed beside each participant’s tree. Symbol shapes are coded by spike protein domain (see legend, center bottom). The frequency of each haplotype detected at each sample timepoint (days post symptom onset) in each participant is indicated to the right of the tree with a scaled black dot. SARS-CoV-2 RNA levels (rRT-PCR Ct values; black traces) and serum antibody binding to spike protein (optical density, 450 nm [OD450]; purple traces) are shown above the dot plot for each participant. The positivity cutoff index (COI) of 0.4 for serum antibody binding to spike protein is indicated with a dashed line.

### Selection and autologous antibody responses

To investigate the intra-host selective forces that might drive SARS-CoV-2 genetic evolution in PWH with CD4 counts <200 cells/μL, we cross-referenced spike gene phylogenetic and selection analyses with autologous antibody analysis for each participant. As described previously^19^, serum binding to the ancestral SARS-CoV-2 spike remained undetectable in many of these individuals through the first 4 weeks of infection (see **Fig. 4**, purple traces, and **Extended Data Fig. 9**), consistent with other recent studies of SARS-CoV-2 humoral immunity in PWH^8,28^. Subsequently, in participants 16-036 and 15-030, serum binding to spike repeatedly exceeded the assay positivity threshold. These delayed but detectable responses were associated with positive selection for RBD mutations linked to immune escape, including presumptive convergent evolution of E484K (see **Fig. 4**, participant 16-036, red pentagons [#7]) and a virus genetic clade defined by L452R (see **Fig. 4**, participant 15-030, red pentagons [#4], top clade on tree). In other PWH with CD4 counts <200 cells/μL, serum spike binding remained undetectable at all timepoints tested. These individuals did not show positive selection within the RBD, but did show positive selection for mutations associated with changes in virus infectivity, including H655Y (see **Fig. 4**, participant 20-017, aqua circle [#5]) and P681H/R (see **Fig. 4**, participant 30-014, aqua circles [#1]). For many selected mutations in PWH with CD4 counts <200 cells/μL, previous studies have provided evidence for antibody evasion and/or increased infectivity **(Extended Data Table 1)**. Interestingly, selected mutations in the NTD that were associated with antibody evasion occurred in both the presence and the absence of detectable serum binding to spike (see **Fig. 4**, participants 20-017 and 15-029, blue hexagons), suggesting that alterations to this region may have important functional impacts beyond humoral immune evasion. Combined with the direct correlation between with early spike diversity and subsequent SARS-CoV-2 RNA shedding duration **(Fig. 2b)**, these findings link intra-host SARS-CoV-2 diversification and adaptive evolution to the persistence of the virus in people with advanced HIV infection.

## Discussion

Defining the extent, kinetics, and evolutionary patterns of SARS-CoV-2 diversification in individuals with immunocompromising conditions is important for understanding both the biology of persistent infections and the emergence of new VOCs. Using specialized sequencing technology to analyze a clinically diverse cohort of PWH and PWOH, we find that permissiveness for SARS-CoV-2 replication in PWH who have low CD4 counts – which was often coupled with uncontrolled plasma HIV viremia – is associated with high levels of SARS-CoV-2 spike genetic diversity just days after COVID-19 symptom onset. Early genetic diversity in these unusual cases is likely necessary for subsequent adaptive evolution, and potentially for intra-host persistence of the infection under changing fitness constraints. Indeed, we find spike gene mutational signatures in individuals with advanced HIV infection that indicate positive selection at sites reported to have important functional roles. Thus, SARS-CoV-2 evolution in these individuals is not solely a product of random diversification through unchecked replication, but instead involves intra-host adaptation that may markedly increase the risk for generation of new SARS-CoV-2 variants.

HIV co-infection likely promotes the intra-host persistence and evolution of SARS-CoV-2 through multiple immune insults. Progressive HIV infection impacts not only the adaptive immune system, but also the type I interferon responses that normally mediate innate antiviral defenses^29,30^. In this regard, we noted that elevated SARS-CoV-2 genetic diversity in PWH with low CD4 T cell counts often preceded the expected onset of adaptive immunity. Innate immune defects in people with advanced and poorly controlled HIV infection could have contributed to this early SARS-CoV-2 diversification by permitting increased early replication and/or a relaxed transmission bottleneck^31,32^. Subsequently, HIV-induced defects in CD4-T-cell-dependent adaptive immunity likely compounded these issues. Replicating HIV may preferentially infect activated, antigen-responsive CD4 T cells^33^, and in progressive disease may also interfere with CD4 T cell help for other lymphocytes by disrupting lymph node architecture^34^. In this setting, weak and delayed humoral immunity to spike risks selecting antibody-escape mutants from a diverse variant pool. It is important to note that antiretroviral therapy (ART) may counteract innate and adaptive immune defects associated with uncontrolled HIV replication^35–38^, and that previous studies have documented similar magnitude and kinetics of SARS-CoV-2-specific immune responses between PWH receiving ART and PWOH^39,40^. Therefore, while our use of specialized sequencing technology was important for describing high intra-host SARS-CoV-2 diversity in this study, our striking findings likely also reflect the unique biology of advanced, uncontrolled HIV infection.

Our findings in this study have several limitations. First, although the HT-SGS technology we used here combines high accuracy with long reads and relatively deep single-molecule sampling, our sequencing was limited to a portion of the virus genome at one anatomic site. Therefore, our results do not reflect the roles of mutations outside spike, differences in virus sequences between tissues, or cumulative virus genetic diversity throughout the body^41,42^. Second, because our study relied on natural infections, we are unable to determine the timing or sources of SARS-CoV-2 transmission to our study participants. We thus cannot rule out that delayed symptom onset and/or transmission of multiple, closely related founders contributed to elevated SARS-CoV-2 diversity detected at early timepoints in some PWH. Limitations on anatomic sampling and immune analysis leave open questions about the relative importance of humoral immune pressure, selection for intra-host transmissibility, and other evolutionary drivers in this setting that may be addressed in future studies using model systems with structural and functional characterization of variant sequences.

Finally, our results do not address how current Omicron subvariants might evolve in people with advanced HIV infection after prior SARS-CoV-2 infection or vaccination, nor do they establish the transmissibility between people of variants identified within each person. Further studies will be needed to understand whether intra-host SARS-CoV-2 variants arising in PWH and those arising in people with other immunocompromising conditions differ in their potential to escape pre-existing immunity in immunocompetent individuals.

Despite these limitations, our results in this study demonstrate the tremendous differences in intra-host SARS-CoV-2 genetic diversity and evolution between people with advanced, poorly controlled HIV infection and those with controlled infection or without HIV infection. The potential emergence of new pandemic virus variants in PWH who are not receiving effective ART remains highly concerning. This concern could be mitigated through active or passive immunizations that provide sufficient early protection to limit virus genetic diversification in this setting. However, our results also emphasize that efforts to control SARS-CoV-2 and potentially other viruses will benefit from addressing remaining gaps in the global approach to HIV infection.

## Methods

### Study Participants

Recruitment of study participants was performed in compliance with relevant ethical regulations. Participants provided informed consent before study.

The hospitalized cohort was enrolled from 20 sentinel surveillance hospitals in 8 of the 9 South African provinces. Cohort enrollments were limited to individuals aged ≥18 years who were living within a 50-kilometer radius of the respective hospitals and who had laboratory-confirmed, symptomatic COVID-19 within 5 days of diagnosis. All cohort participants underwent a combined nasopharyngeal/oropharyngeal swab at enrollment and every second day thereafter until cessation of SARS-CoV-2 shedding, as defined by 2 consecutive negative swabs. Serum specimens were collected at enrollment and days 7, 14 and 21 post symptom onset^19^. We selected individuals from the hospitalized cohort for inclusion in the present study if they had SARS-CoV-2 N gene rRT-PCR Ct ≤30 on their initial samples and at least 3 positive samples. Demographic and clinical information was collected using standardized case report forms at enrollment; daily while in hospital; and at discharge from hospital, when shedding stopped or when the individual died.

The case-ascertained household transmission study, which included the outpatient cohort, took place in Klerksdorp (North West Province) and Soweto (Gauteng Province), South Africa. Screening of index cases occurred at three clinics in Klerksdorp from October 2020 to June 2021, and at five clinics in Soweto from October 2020 to September 2021. Individuals aged ≥18 years with COVID-19-compatible symptoms starting ≤5 days before presentation were screened for SARS-CoV-2 on nasopharyngeal swabs at primary health clinics. Households of individuals positive for SARS-CoV-2 were enrolled if the index case symptoms started within 7 days before household enrollment, if no other household members reported symptoms 14 days prior to household enrollment, and if there were ≥2 additional household members of whom ≥70% provided consent for study. Households were followed for six weeks, with nasal swabs collected three times per week and serum samples collected at baseline and at the end of the follow-up period^20^. From the outpatient cohort, we selected individuals for the present study who tested positive for SARS-CoV-2 with SARS-CoV-2 N gene rRT-PCR Ct ≤35. Household, demographic, and clinical information was collected in this cohort at enrollment. Information about symptoms and healthcare-seeking behavior were collected at thrice weekly follow-up visits using Research Electronic Data Capture (REDCap) databases on electronic tablets.

In the hospitalized cohort, HIV testing was conducted as part of clinical management. If an HIV diagnostic test result was not obtained during a participant’s hospitalization, a prior documented positive result or evidence in hospital records of treatment with ART was considered to indicate HIV positivity. A documented negative HIV diagnostic test result within 6 months of study was considered to indicate HIV negativity. Participants with unknown HIV status or with negative HIV diagnostic test results older than 6 months were offered voluntary counselling and testing by rapid enzyme-linked immunosorbent assay (ELISA).

In the outpatient cohort, rapid HIV testing was offered for individuals with unknown HIV status, or for whom a documented negative HIV result was not available within the previous 6 months. For one participant aged 0.5 years whose mother had a documented HIV negative status during pregnancy, HIV status was considered negative. For individuals who did not agree to rapid testing, but did consent to HIV testing, residual serum was tested for HIV antibodies by ELISA at the National Institute for Communicable Diseases (NICD). For PWH, data on CD4 counts and plasma HIV RNA levels within 6 months of enrollment were collected from medical records. If these data were not available, samples were collected and plasma HIV RNA levels tested by quantitative rRT-PCR (Roche Cobas Ampliprep/Cobas Taqman HIV-1 test, Roche Diagnostics, Mannheim, Germany) at the NICD.

For the hospitalized cohort, ethical clearance was obtained through the University of the Witwatersrand health research ethics committees (HREC) (Medical) (M160667); Stellenbosch University HREC (15206); University of Pretoria HREC (256/2020), and University of the Free State HREC (HSD2020/0625). For the outpatient cohort, clearance was obtained from the University of the Witwatersrand HREC (M2008114). Participants in the outpatient cohort received a $3.00 grocery store voucher at each follow-up visit to compensate for time required for specimen collection and interview.

### Detection of SARS-CoV-2 RNA in swab specimens

Upper respiratory specimens (combined nasopharyngeal and oropharyngeal specimens in the hospitalized cohort, nasopharyngeal specimens for screening of index cases in the outpatient cohort, and nasal specimens for household follow-up in the outpatient cohort) were collected by trained study nurses using nylon flocked swabs and transported in viral or universal transport medium to the NICD for further testing. Total nucleic acids were extracted from 200 µl of each sample using the DNA/Viral NA Small Volume v2.0 extraction kit (Roche Diagnostics, Mannheim, Germany) and an automated extractor MagNA Pure 96. Detection of SARS-CoV-2 nucleic acid from specimens was performed using the Allplex™ 2019-nCoV assay (Seegene, Seoul, South Korea) with rRT-PCR. Specimens were considered positive for SARS-CoV-2 if the Ct value was <40 for any of the E, RdRp and N SARS-CoV-2 gene targets.

### HT-SGS

Aliquots of swab samples stored at −80°C were thawed at room temperature and centrifuged briefly before RNA extraction. Virus RNA was extracted with a magnetic bead-based RNA extraction kit (RNAdvance Viral Reagent kit, Beckman Coulter, C63510) on an epMotion® 5073t liquid handler (Eppendorf, 5073000345). Extracted RNA was immediately reverse-transcribed. Procedures for reverse-transcription and for purification, quantification, and PCR amplification of complementary DNA (cDNA) were as previously described^18^. Reverse-transcription was performed using SuperScript IV Reverse Transcriptase (ThermoFisher Scientific, 18090010) according to the manufacturer’s instructions. The reverse-transcription primer consisted of an outer reverse primer binding site for PCR, an 8-base unique molecular identifier (UMI) of randomly incorporated bases, and a gene-specific target region (CCGCTCCGTCCGACGACTCACTATACCCGCGTGGCCTCCTGAATTATNNNNNNNNCGTTGCAGTAGCGCGAACAA). After reverse-transcription, cDNA was treated with proteinase K (Sigma-Aldrich, 3115828001) for 25 min at 55°C with shaking at 1000 rpm to digest residual protein, followed by purification using RNAClean XP bead suspension (A63987, Beckman Coulter) at a bead:cDNA volume ratio of 2.2:1. The cDNA copy number in a small aliquot of each sample was measured on a QIAcuity digital PCR (dPCR) system (Qiagen) using forward primer ACGTGGTGTTTATTACCCTGACA, reverse primer TTGGTCCCAGAGACATGTATAGC, and hydrolysis probe 5′-/56-FAM/FAM TTTCCAATGTTACTTGGTTCCA/3BHQ_1/-3′ (synthesized by IDT). Cycling conditions were as follows: initial denaturation at 95°C for 2 min, followed by 45 cycles of 95°C for 15 sec and 53°C for 1 min. After dPCR quantification, full-length, UMI-tagged spike gene cDNA was amplified using the Advantage 2 PCR kit (Takara Bio, 639206) with forward primer TTCGCATGGTGGACAGCCTTTGTT and reverse primer CCGCTCCGTCCGACGACTCACTATA under the following thermocycling conditions: initial denaturation at 95°C for 1 min; 32 cycles of 95°C for 10 sec, 64°C for 30 sec, and 68°C for 5 min; and final extension at 68°C for 10 min. PCR reagents concentrations were as follows: 800 nM forward and reverse primers, 400 μM dNTP, 1X Advantage 2 Buffer, and 2X of Advantage 2 Polymerase Mix. For long-read sequencing, amplified DNA products of length 4.3-kilobases (encompassing the entire 3.8-kilobase spike gene) were incorporated into sequencing libraries using the SMRTbell Express Template Prep Kit 2.0 (100-938-900, Pacific Biosciences) and Barcoded Overhang Adapter kit 8A and 8B (101-628-400 and 101-628-500, Pacific Biosciences) for multiplexing targeted sequencing. Libraries were processed through primer annealing and polymerase binding using the Sequel II Binding Kit 2.0 (101-842-900, Pacific Biosciences), and then sequenced on a Sequel II system (Pacific Biosciences) with a 20-hour movie time under circular consensus sequencing (CCS) mode.

### SGS calling

Circular consensus sequences (CCS) were generated from SMRT sequencing data with minimum predicted accuracy of 0.99 and a minimum of 3 passes in Pacific Biosciences SMRT Link (v11.0.0.146107)^43^. CCS reads were demultiplexed using Pacific Biosciences barcode demultiplexer (lima) to identify barcode sequences. The resulting FASTA files were reoriented into the 5’-3’ direction using the vsearch —orient command in vsearch (v2.21.1). Cutadapt (v4.1) was used to trim forward and reverse primer sequences. Length filtering was performed to remove reads shorter than 2800 nt or longer than 4000 nt. Remaining reads were then binned by their 8-base UMI sequences. For each bin, reads were clustered with vsearch —cluster_fast based on 99% sequence identity. Only bins that yielded a single, predominant cluster (i.e., where the largest cluster was (1) inclusive of at least half of the bin’s reads and (2) at least twice as large as the second largest cluster) with at least 10 CCS reads were kept. The cluster consensus sequence generated by the vsearch —cluster_fast was then used as a reference to map the cluster’s reads with minimap2 (v2.24). The commands bcftools mpileup −X pacbio-ccs and bcftools consensus were used to determine the final consensus sequence for each bin. Final consensus sequences were used as queries for BLAST nt database searches, and non-SARS-CoV-2 sequences thus identified were discarded.

Putative false UMI bins (spurious bins that arise due to PCR and/or sequencing errors) were identified and removed with a network approach as previously described^18^. Given two distinct bins a and b with read counts n_a_ and n_b_, and assuming n_a_ ≥ n_b_, a and b are connected by an edge if they have edit distance 1 and satisfy the following count criterion: n_a_ ≥ 2n_b_–1. Networks formed as above were resolved using the adjacency method^44^, which iteratively consolidates smaller bins into larger bins that meet the above criteria. As a final filter, a mixture model of bin size was iteratively optimized using exponential and Gaussian distributions representing false and real bins, respectively. Bins with posterior probability Prob(false) >0.5 were discarded and the remaining bins were used as final SGS. The full HT-SGS data processing pipeline used is available at https://github.com/niaid/UMI-pacbio-pipeline/releases.

### Variant and haplotype calling

Despite high CCS read accuracy and UMI-based error correction, sample reverse-transcription errors and other rare errors nonetheless persist in processed HT-SGS datasets. To address such errors, variant calling was performed using a model describing technical error rates. Given a reverse-transcription (RT) error rate R = 1 × 10^-4^, a target insert length L, and number of recovered SGS sequences N, the probability of observing a technical variant with at least *c* occurrences in the sample was expressed as P(C≥*c*) = 1 – Binom_CDF_(*c* | N, R)^L^. To determine a cutoff for variant calling, the smallest value *c_v_* for which P(C≥*c_v_*) <0.01 was determined, and the minimum number of occurrences to call a variant was set as *c_v_* + 1. Indels were handled separately; at least three identical occurrences of an indel were criteria for inclusion as a real variant. Variants in each sample not meeting these criteria were reverted to the consensus of all SGS for that sample. This variant calling approach was implemented with a custom Python script that is available at https://github.com/niaid/UMI-pacbio-pipeline/releases. Among SGS subjected to variant calling, each unique combination of mutations within the individual was considered as one haplotype. To avoid inaccurate findings arising from any residual erroneous sequences, only SGS representing haplotypes that were detected at least 2 times in each sample were included in downstream analysis.

### Normalized Shannon entropy

Normalized Shannon Entropy (H_norm_) for a group of aligned sequences was calculated as the entropy for those sequences (H) divided by the maximum possible entropy for that number of sequences (H_max_), 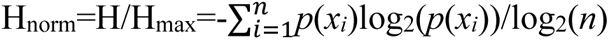.

### Average pairwise genetic distance

Average pairwise distance for each group of sequences was calculated as the total number of mutations (point mutations and/or indels) between each pair of sequences divided by the number of pairs in that group.

### Jensen-Shannon distance

Within each participant, spike sequence population dissimilarity (distance) was analyzed for all possible sample pairs. The Jensen-Shannon calculation was performed for each sample pair using the haplotype frequency distributions for the samples and the “jensenshannon” method in *SciPy* (v 1.8.1).

### Estimation of mutations below detection limit

We anticipated that some real variant sequences might be inadvertently removed by our data analysis process due to sampling depth limitations. To estimate the number of uncalled biological mutations (MU) in each sample after analysis, we considered the total number of mutations in the sample before variant calling (M0), the number of mutations that exceeded the variant calling threshold and were therefore interpreted as real (M1), and an estimate of the number of technical mutations (e.g., RT errors) expected in the sample (M2; based on an assumption of 1 x 10^-4^ error/base). M0 and M1 were counted directly from an alignment of all SGS for the sample, and M2 was computed as the upper 99% CI of a Binomial distribution with N as the total number of nucleotides in the alignment and *p* = 1 x 10^-4^, as computed via SciPy (v 1.8.1). MU was then calculated as MU = M0 – M1 – M2 and restricted to be zero or greater. We considered values of MU >0 to imply the presence of real mutations that went uncalled after the variant and haplotype calling process.

### Phylogenetic inferences

Recombinant sequences were identified using 3SEQ^45^ (v 1.8.0) using “full run” mode on each participant’s haplotypes and a threshold of *p* <0.05. Recombinant sequences thus identified were excluded from initial phylogenetic models and analyses of positive selection.

Phylogenetic relationships were inferred within each participant using all non-recombinant haplotype sequences for that participant and with Wuhan-Hu-1 spike (GenBank Accession NC_045512.2, nucleotide coordinates 21563-25384) included as an outgroup. To account for indels when performing phylogenetic analyses, a binary matrix was generated using 2matrix^46^, which encoded for the presence or absence of indels in each haplotype. This matrix was computed using a curated combined alignment of all haplotypes across all participants. Trees were then constructed using iqtree^47^ (v 1.6.12) with a partitioned model. Nucleotide sequences were analyzed using an HKY model, while the indel matrix was analyzed using a JC2 morphological model with enforced transition rates of 0.99 (indel acquisition) and 0.01 (indel reversion). Maximum-likelihood trees were computed, and support values were obtained through ultra-fast bootstrapping with 1000 iterations.

The presence of multiple founder sequences was assessed using TreeCluster^23^. Clusters were formed using the “single linkage” mode of TreeCluster with a phylogenetic distance threshold of 0.0015, which corresponds to >5 mutations within the 3822 nt coding sequence of spike. For participants in whom multiple clusters were detected, each cluster was re-processed with the phylogenetic method described above to yield subtrees corresponding to each founder; recombinant sequences identified earlier were added to the associated cluster most divergent from Wuhan-Hu-1. The subtrees were concatenated into a single tree with a custom Python script, and then branch lengths were reoptimized with iqtree.

### Testing for selection

Testing for positive selection was performed initially with FUBAR^27^ using trees of participant haplotypes computed as described above. Outlying sequences in each participant with less than 80% amino acid homology to other sequences were excluded before analysis, thereby removing most premature stop codons, large deletions, and frameshifts. For ω equal to the relative rate of nonsynonymous over synonymous mutations at a site, we considered sites with Prob(ω >1) >0.9 to be under possible positive selection. To confirm positive selection for these sites, we examined frequencies of associated non-synonymous mutations in the participant over time. Frequency changes were assessed relative to the participant’s first sample and considered the summed frequencies of all haplotypes containing a given mutation. Mutations with a frequency increase of at least 0.20 in the participant and Prob(ω >1) >0.9 via FUBAR were called as under positive selection.

### Short-read WGS data

Short-read data from selected samples from both the hospitalized and outpatient cohorts were obtained with the Ion Torrent Genexus platform using AmpliSeq for SARS-CoV-2, following the ARTIC SARS-CoV-2 sequencing protocol, as previously described^19,20^. Data were processed with a unified pipeline in CLC Genomics Workbench v22.0.3. This pipeline included quality filtering, read trimming, mapping to the Wuhan-Hu-1 reference (Hu-1, GenBank Accession NC_045512.2), local realignment, consensus calling, and variant calling. Variant calling was performed with a minimum coverage of 10, minimum variant count of 2, and minimum variant frequency of 1%.

### Measurement of spike antibody binding responses

In both the hospitalized and outpatient cohorts, antibodies against the SARS-CoV-2 spike protein were detected using an enzyme-linked immunosorbent assay (ELISA) as previously described^48^. Recombinant trimeric spike protein was coated onto 96-well, high-binding plates at a concentration of 2 μg/ml and incubated overnight at 4°C. Subsequently, the plates were washed and blocked using a blocking buffer containing 5% skimmed milk powder, 0.05% Tween 20, and 1× PBS, followed by incubation at 37°C for 1–2 hours. Serum samples diluted 1:100 and control antibodies (positive: CR3022 and negative: Palivizumab) diluted to 10 μg/mL were added to the plates, followed by a one-hour incubation at 37°C. Next, an anti-human horseradish peroxidase-conjugated antibody was added and incubated for another one hour at 37°C. To visualize the antibody binding, a OneStep TMB substrate (Thermo Fisher Scientific, USA) was added and allowed to develop for 5 min at room temperature. The reaction was stopped by adding 1 M H_2_SO_4_ stop solution. The absorbance at 450 nm was measured, and specimens with optical density (OD) >0.4 were considered positive for anti-spike antibodies.

### Solvent accessible surface area (SASA) in the spike trimer

Solvent accessible surface area was assessed via NACCESS^25^ on an atomistic structure model of the spike trimer generated using YASARA (htttp://www.yasara.org) on a D614G structure template (PDB: 7KRQ)^26^. The accessibility of each residue was computed as the mean of the 3 accessibilities of the residue over the 3 protomers in the trimer.

### Comparison of spike mutations with public data

Publicly available SARS-CoV-2 spike nucleotide sequences were downloaded from the NCBI Virus database (https://www.ncbi.nlm.nih.gov/labs/virus/vssi/#/), filtering for the South Africa region and collection dates between April and October 2020. The spike sequences were extracted from the downloaded sequences and aligned to Hu-1, and the number of relative point mutations was counted for each sequence. This process was repeated for each intra-host SGS from within the collection period.

### Analysis of the intra-host spike sequences with Nextclade

All intra-host sequences were uploaded to Nextclade web (https://clades.nextstrain.org). The results were downloaded in the nextclade.tsv file and the Nextclade_pango lineage field was extracted for each of the input sequences. The relative frequency of each assigned Nextclade Pango lineage per host was used to plot the frequencies of intra-host Pango lineages.

### Statistical analyses

GraphPad Prism v.9.3.1 and MATLAB 2022a were used for statistical analyses. Specific statistical tests used in each analysis are presented in the corresponding figure legend. The significance of single comparisons in multiple groups was assessed by one-way ANOVA with multiple comparisons using the Kruskal-Wallis test (unpaired or unmatched groups) or Friedman test (paired groups) and Dunn’s multiple comparisons test. The nonparametric Spearman’s test (two-tailed) and simple linear regression were used for correlation analyses. Modeling of the primary major haplotype frequency was performed with logistic regression via the method of iteratively reweighted least squares.

## Acknowledgments

We gratefully acknowledge the participants in this study. We thank Sibongile Walaza for assistance with the clinical studies. This work was supported by the NIH Intramural Research Program (Vaccine Research Center), and by the Wellcome Trust (grant 221003/Z/20/Z) in collaboration with the Foreign, Commonwealth and Development Office, United Kingdom, the US Centers for Disease Control and Prevention (co-operative agreement 6 U01IP00104804-02), as well as the National Institute for Communicable Diseases, a division of the National Health Laboratory Service, South Africa.

## Author Contributions

Conceptualization – SHK, JNB, AVG, CC, EAB

Sample acquisition and primary sample testing – JNB, SM, JK, DA, DK, NM, LL, JE, ST, NW, AVG, CC

Management of primary study – JNB, SM, JK, NM, LL, ST, NW, AVG, CC HT-SGS of SARS-CoV-2 spike – SHK, ML

Bioinformatic analysis – PR, FB

Phylogenetic and Evolutionary analysis – PR, FB, VGC Short-read whole-genome sequencing – DA, JNB, DK, NW

Solvent accessible surface area (SASA) analysis – TB, RR, PDK Resources – AVG, CC, EAB

Manuscript, original draft – SHK, PR, FB, EAB Manuscript, review and editing – all authors Supervision – AVG, CC, EAB

## Additional information

**Supplementary Information is available for this paper**

## Data Availability

Long-read sequencing data that support the findings of this study have been deposited to the Sequence Read Archive under PRJNA1055920. (https://dataview.ncbi.nlm.nih.gov/object/PRJNA1055920?reviewer=3hd9f3oakpf6vindnbbtsqnkpm)

## Code Availability

UMI-pacbio-pipeline v.1.1 was used to generate single-genome sequences and call haplotypes. This pipeline is available at https://github.com/niaid/UMI-pacbio-pipeline. Additional code used to generate data supporting the findings of this study were deposited at https://github.com/niaid/UMI-pacbio-pipeline/releases/tag/1.1-4c76b04.

## Extended Data

**Extended Data Fig. 1.**
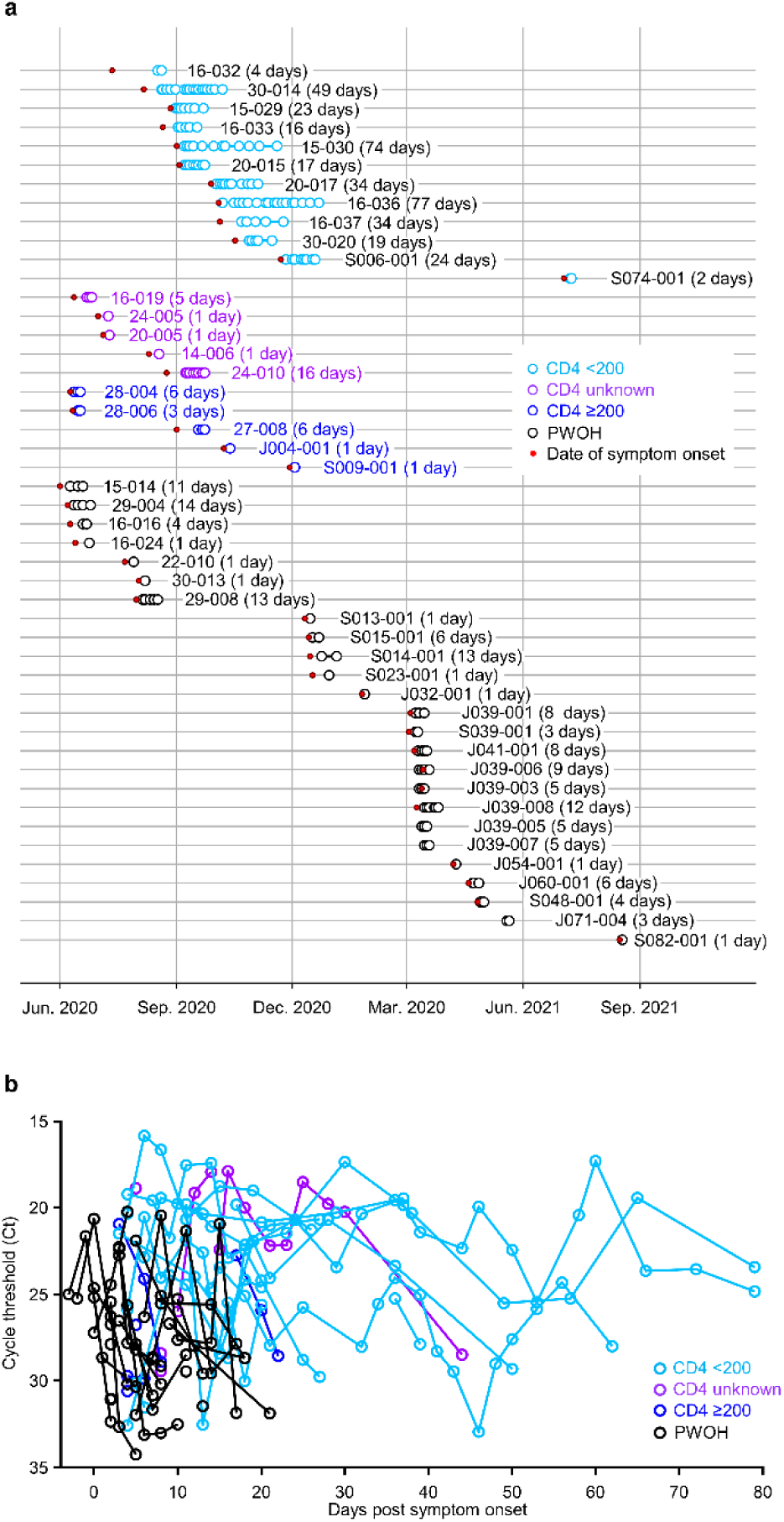
Respiratory sampling in PWH and PWOH. (a) Sampling timelines for all participants. Red dots indicate dates of symptom onset. Next to each participant identifier, the time between the first and last sequenced sample (i.e., the sequenced sampling duration) is indicated. (b) SARS-CoV-2 RNA levels (rRT-PCR Ct values) over time.

**Extended Data Fig. 2.**
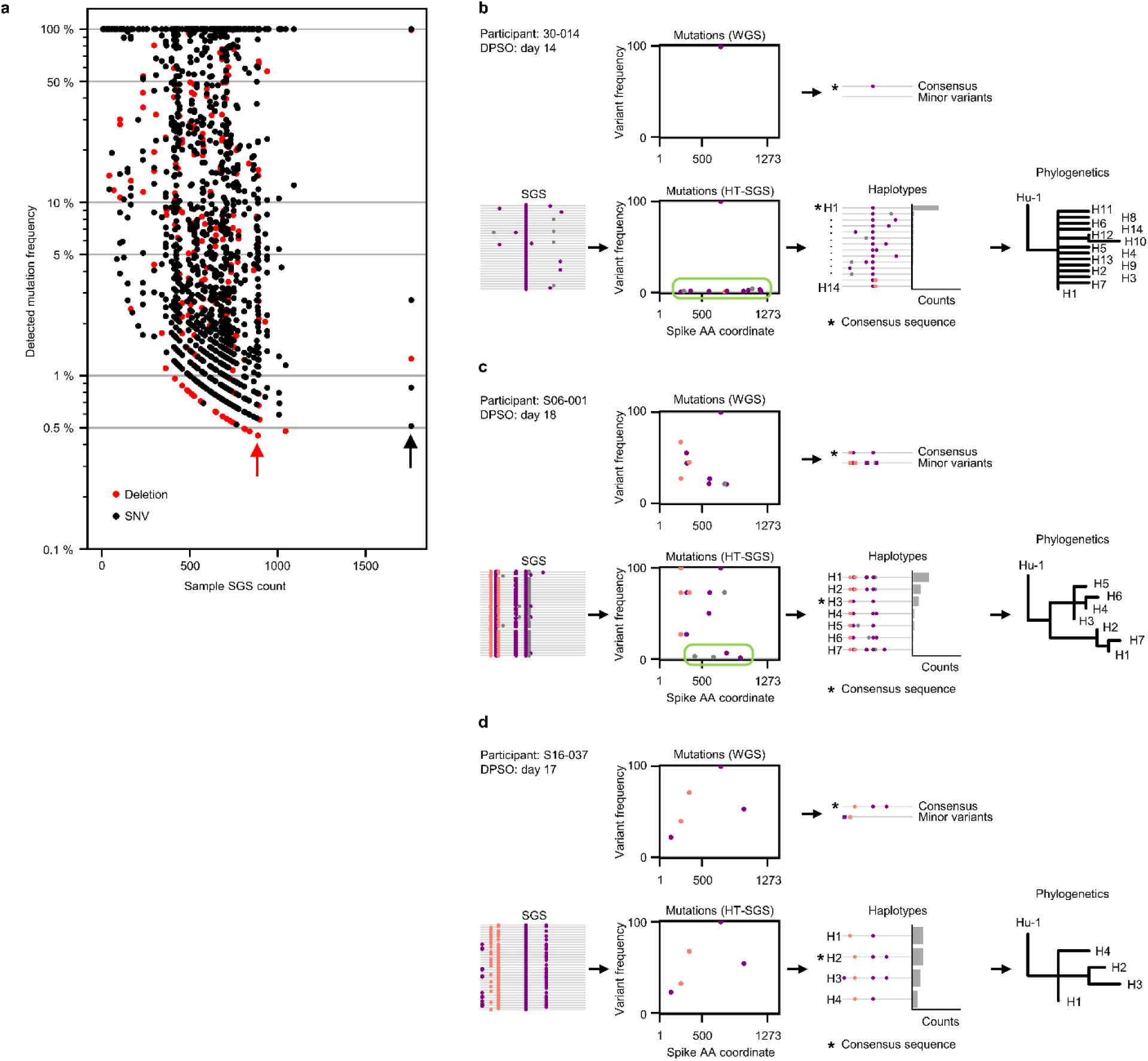
HT-SGS vs. standard whole-genome sequencing (WGS). (a) Frequencies of intra-host mutations detected by HT-SGS as a function SGS numbers for all samples sequenced. Marker color indicates mutation type (red, deletion; black, SNV). The lowest-frequency mutation of each type detected among all samples in the study is indicated by an arrow. (b-d) Comparisons of results obtained from standard, short-read-based WGS (upper half of each panel) and HT-SGS (lower half of each panel) for three selected samples. Participant identifiers and sample timepoints (in days post symptom onset [DPSO]) are indicated. For WGS, consensus sequences and minor variant mutations are indicated. For HT-SGS, all detected haplotypes and as well as phylogenies that relate haplotypes to one another are shown.

**Extended Data Fig. 3.**
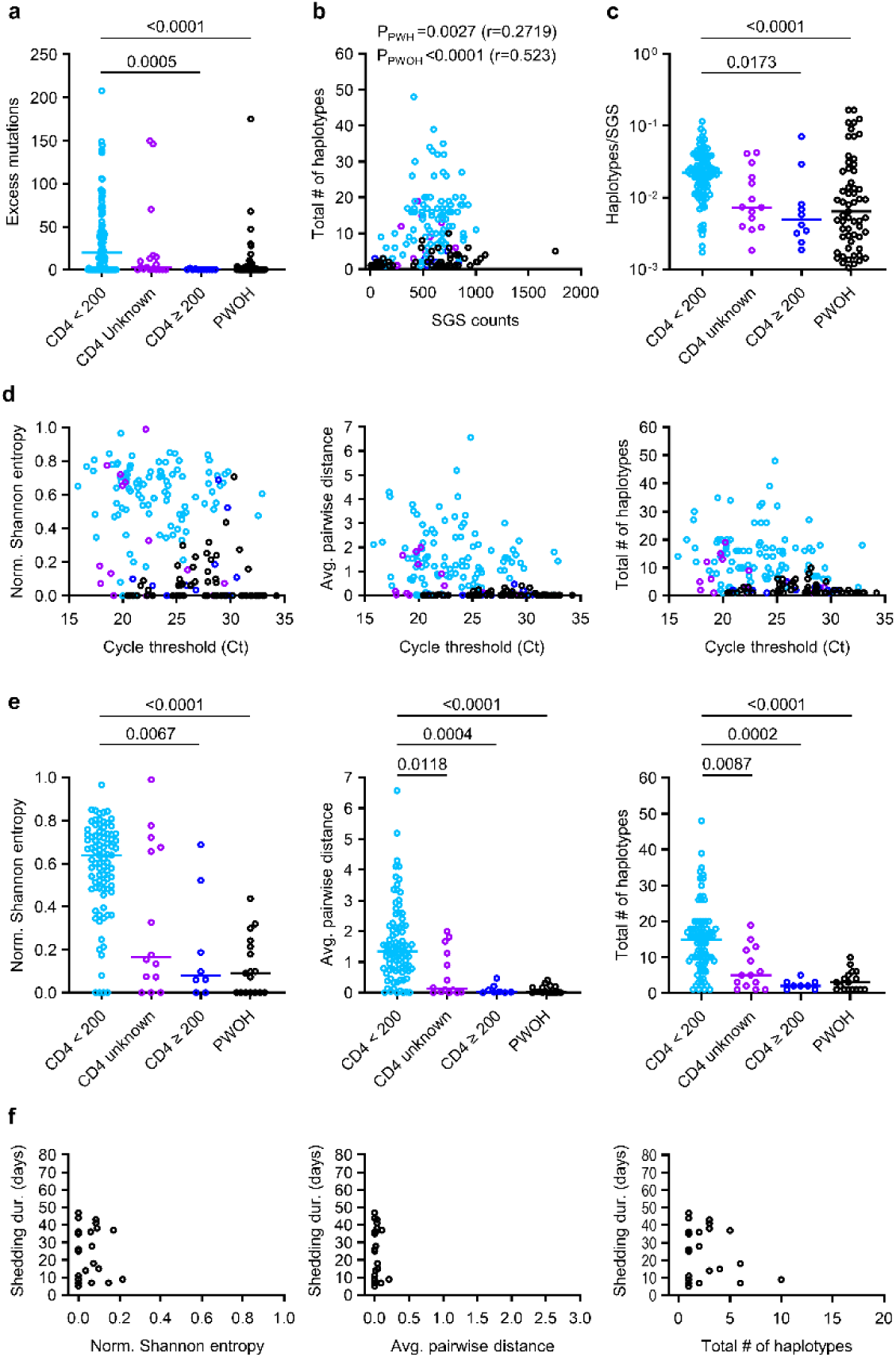
Correlation among intra-host spike diversity, SARS-CoV-2 RNA levels, and SARS-CoV-2 RNA shedding duration in PWH and PWOH. (a) Excess low-frequency mutations below the detection limit for each sample sequenced in the study. Excess mutations are defined as the estimated number of uncalled real mutations remaining in a sample after considering the number of called real variants and an estimated number of technical errors, and were calculated as described in Methods. Statistical significance was assessed by one-way ANOVA with multiple comparisons (Kruskal-Wallis test and Dunn’s multiple comparisons test); *p* values <0.05 are shown. (b) Numbers of haplotypes detected across individual samples as a function of SGS counts. Spearman *p* values for correlations between haplotype numbers and SGS counts are shown for PWH and PWOH. (c) Ratios of haplotypes identified per SGS. Statistical significance was assessed by one-way ANOVA with multiple comparisons (Kruskal-Wallis test and Dunn’s multiple comparisons test); *p* values <0.05 are shown. (d) Correlations between measurements of spike diversity and SARS-CoV-2 RNA levels (rRT-PCR Ct values). (e) Comparison of spike genetic diversity among PWH subgroups and PWOH in the hospitalized cohort. Individual samples from longitudinal sample sets in each person are represented by separate datapoints. Statistical significance was assessed by one-way ANOVA with multiple comparisons (Kruskal-Wallis test and Dunn’s multiple comparisons test); *p* values <0.05 are shown. (f) Correlations between measurements of spike diversity at the first sample timepoint and SARS-CoV-2 RNA shedding duration in PWOH.

**Extended Data Fig. 4.**
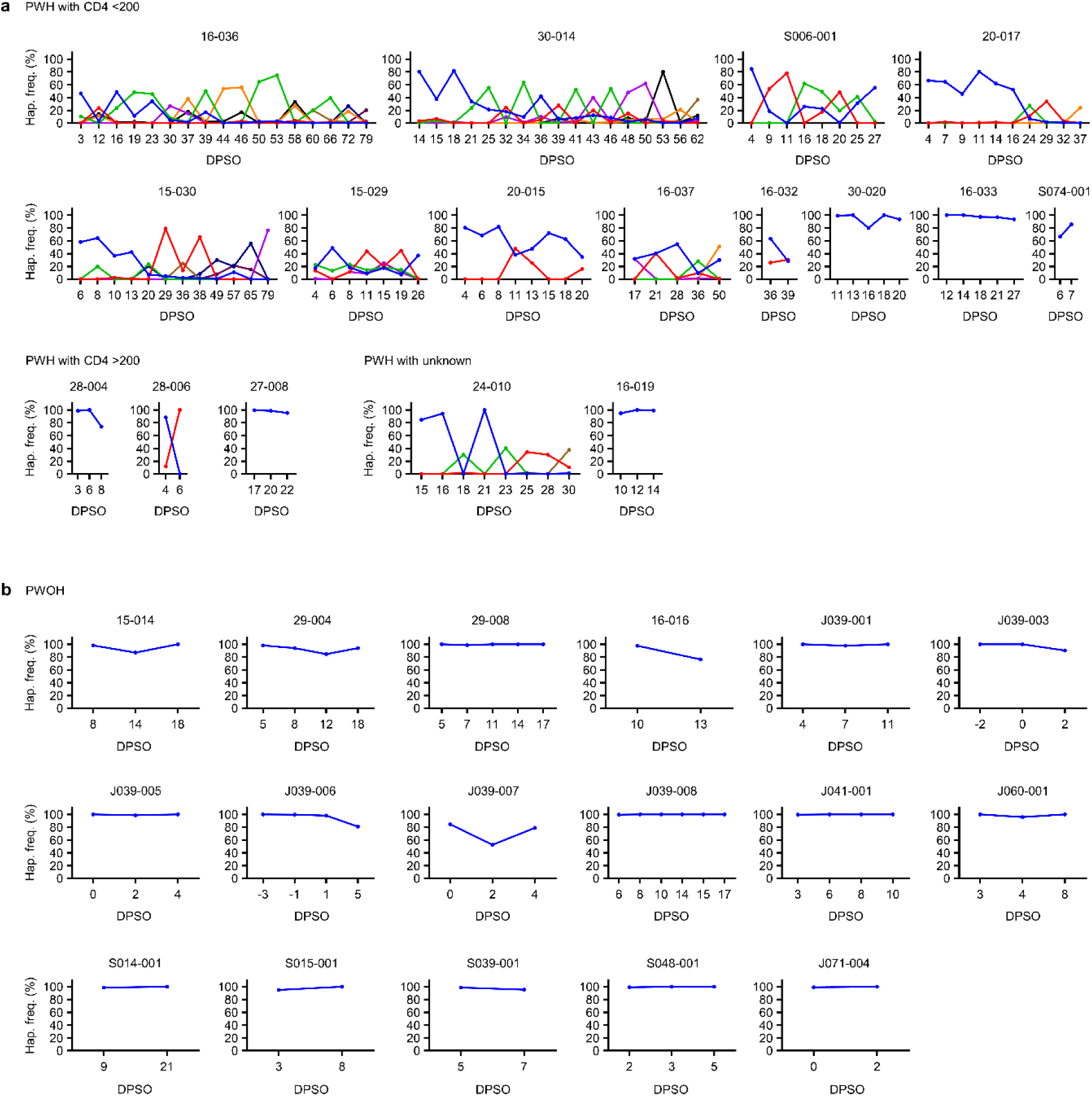
Frequencies of abundant spike haplotypes in each PWH or PWOH over time. The frequency of the haplotype that was most abundant at the first timepoint in each participant is indicated in blue, with time indicated on the x-axis as DPSO. Frequencies of other haplotypes that were most abundant in 1 or more other timepoints are also shown. Results are shown for PWH (a) and PWOH (b).

**Extended Data Fig. 5.**
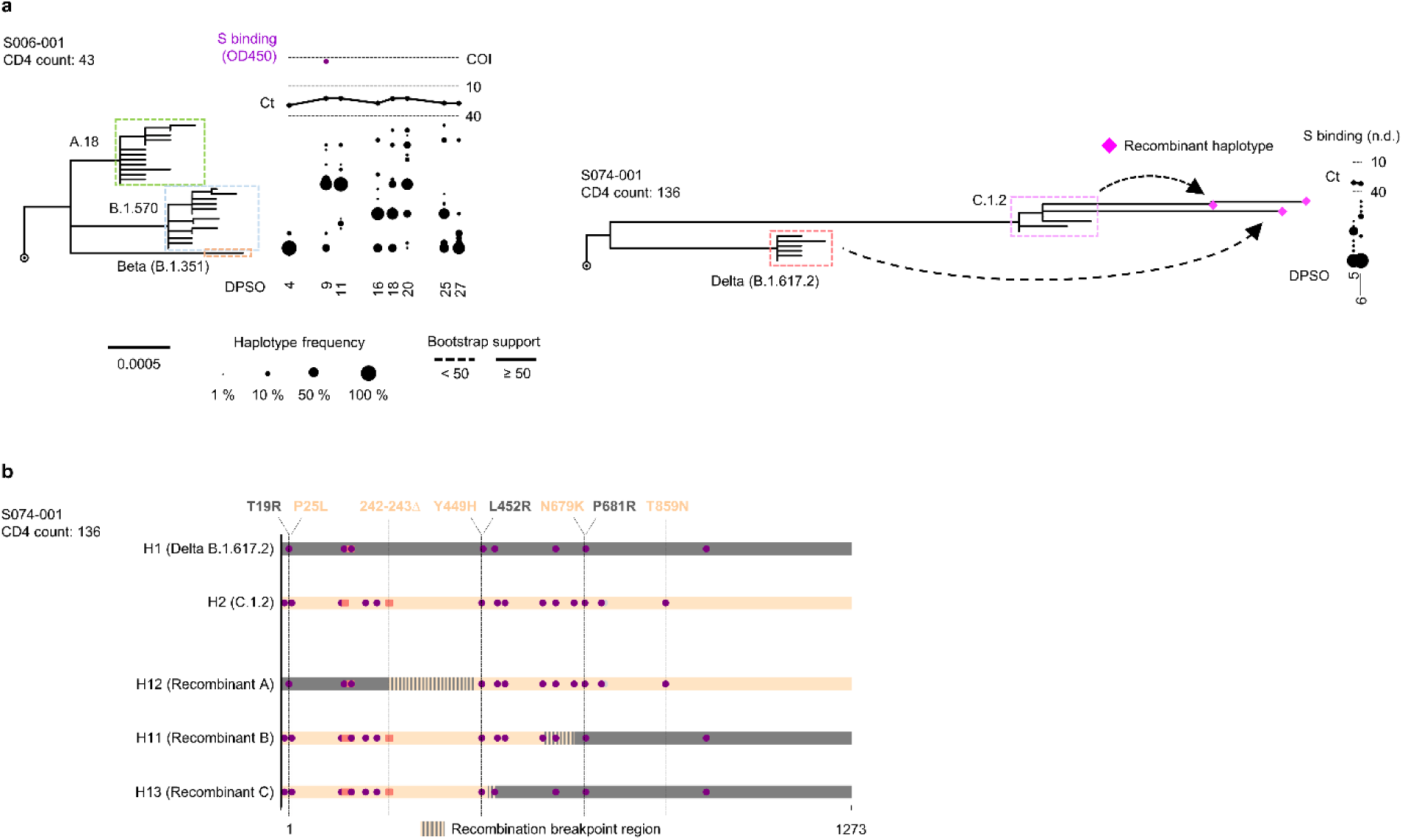
Multiple founder sequences and intra-host recombination in some PWH with CD4 counts <200 cells/μL. (a) Maximum-likelihood phylogenetic trees rooted on Hu-1 for all haplotypes from the two PWH with CD4 counts <200 cells/μL who were infected with multiple founders, as determined via single-linkage phylogenetic clustering with TreeCluster^23^. Sequences from participant S074-001 that were detected as intra-host recombinants using 3SEQ^45^ are indicated as pink diamonds. Clades with bootstrap support less than 50% are indicated with dashed lines. The frequency of each haplotype detected at each sample timepoint in each participant is indicated to the right of the tree with a scaled black dot. SARS-CoV-2 RNA levels (rRT-PCR Ct values; black traces) and serum antibody binding to spike protein (optical density, 450 nm [OD450]; purple traces; n.d.-no data) are shown above the dot plot for each participant. The positivity cutoff index (COI) of 0.4 for serum antibody binding to spike protein is indicated with a dashed line. (b) Bar-plot of three haplotypes in participant S074-001 that were deemed as intra-host recombinants. Mutations are represented as purple circles (nonsynonymous mutations), grey circles (synonymous mutations), and orange squares (deletions). The color of each bar (dark grey and light orange) represents the presumed origin of the segment; segments with vertical stripes represent the inferred breakpoint regions for each recombinant haplotype.

**Extended Data Fig. 6.**
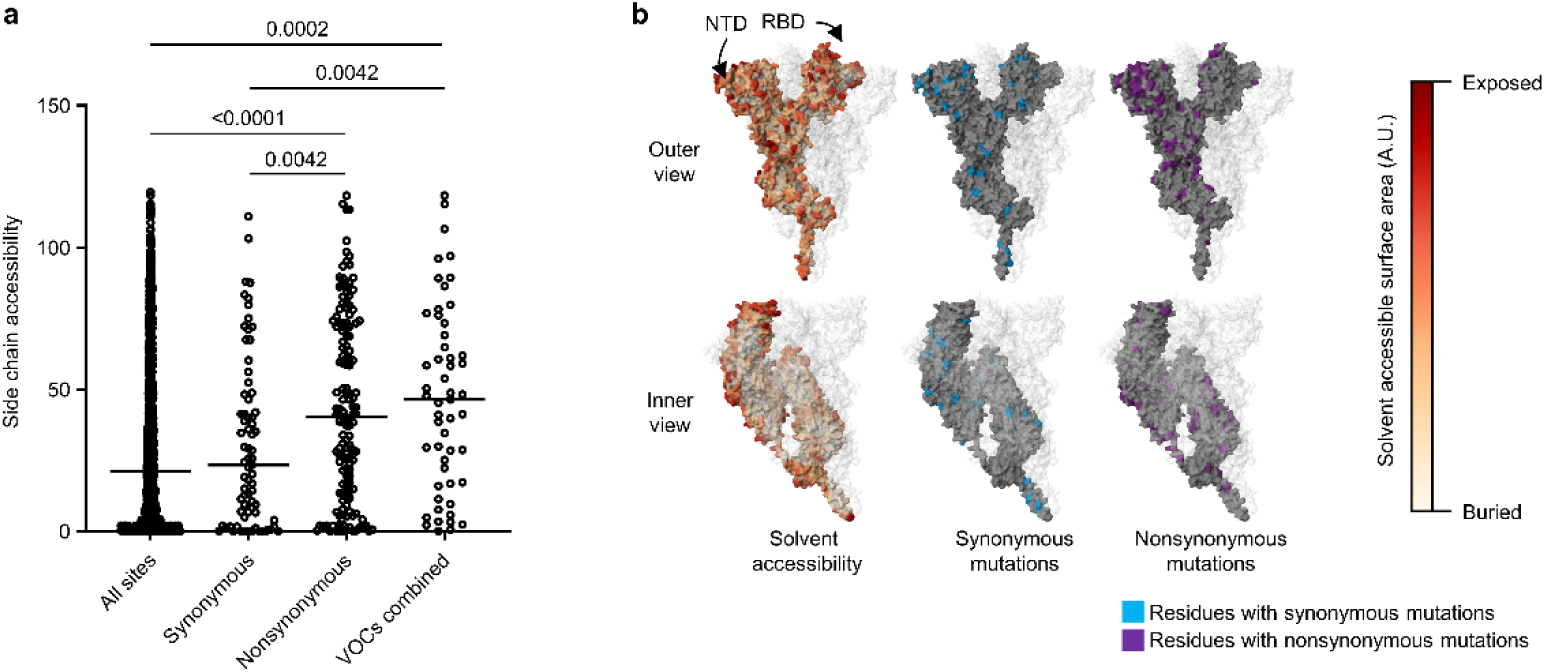
Spike amino acid side chain accessibility in PWH with CD4 counts <200 cells/μL and in VOCs. (a) Measured side chain accessibility of amino acid residues in a structure model of the spike trimer. The “All sites” column refers to all 1273 amino acid residues in the spike protomer. “Synonymous” and “Nonsynonymous” columns refer to residues with intra-host mutations in PWH with CD4 counts <200 cells/μL. The “VOCs combined” column refers to residues that are mutated in Alpha, Beta, Delta, and/or Omicron BA.1 VOCs. Statistical significance was assessed by one-way ANOVA with multiple comparisons (Kruskal-Wallis test and Dunn’s multiple comparisons test); *p* values <0.05 are shown. (b) Structural representation of side chain accessibility (left), synonymous mutations (middle), and nonsynonymous mutations (right) observed in PWH with CD4 counts <200 cells/μL. Results are shown on one spike protomer, with the other two protomers of the trimer made transparent.

**Extended Data Fig. 7.**
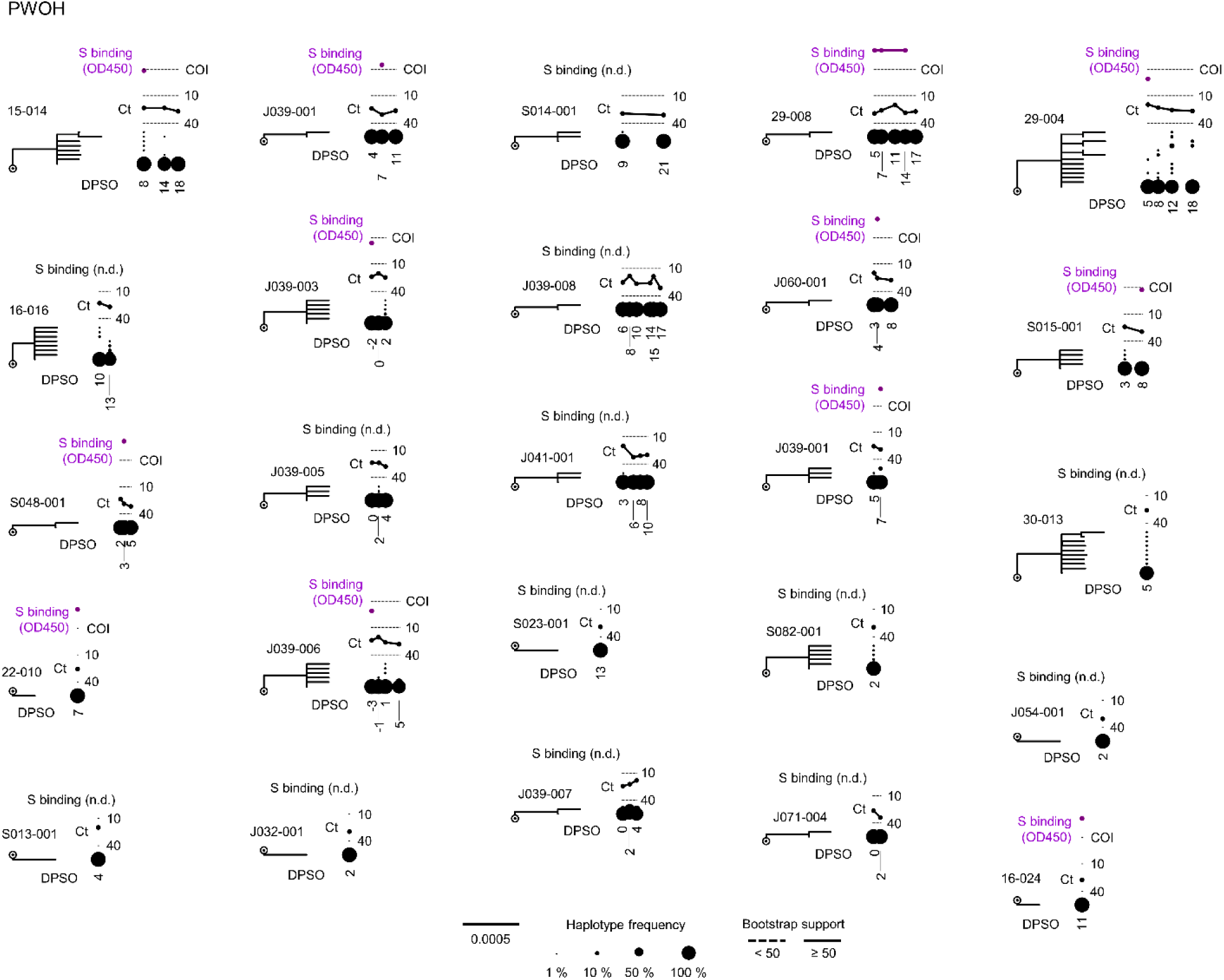
SARS-CoV-2 spike evolution in PWOH. Maximum-likelihood phylogenetic trees rooted on Hu-1 for all spike haplotypes from each PWOH. Clades with bootstrap support less than 50% are indicated with dashed lines. The frequency of each haplotype detected at each sample timepoint (in DPSO) in each participant is indicated to the right of the tree with a scaled black dot. SARS-CoV-2 RNA levels (rRT-PCR Ct values; black traces) and serum antibody binding to spike protein (optical density, 450 nm [OD450]; purple traces; n.d.-no data) are shown above the dot plot for each participant. The positivity cutoff index (COI) of 0.4 for serum antibody binding to spike protein is indicated with a dashed line.

**Extended Data Fig. 8.**
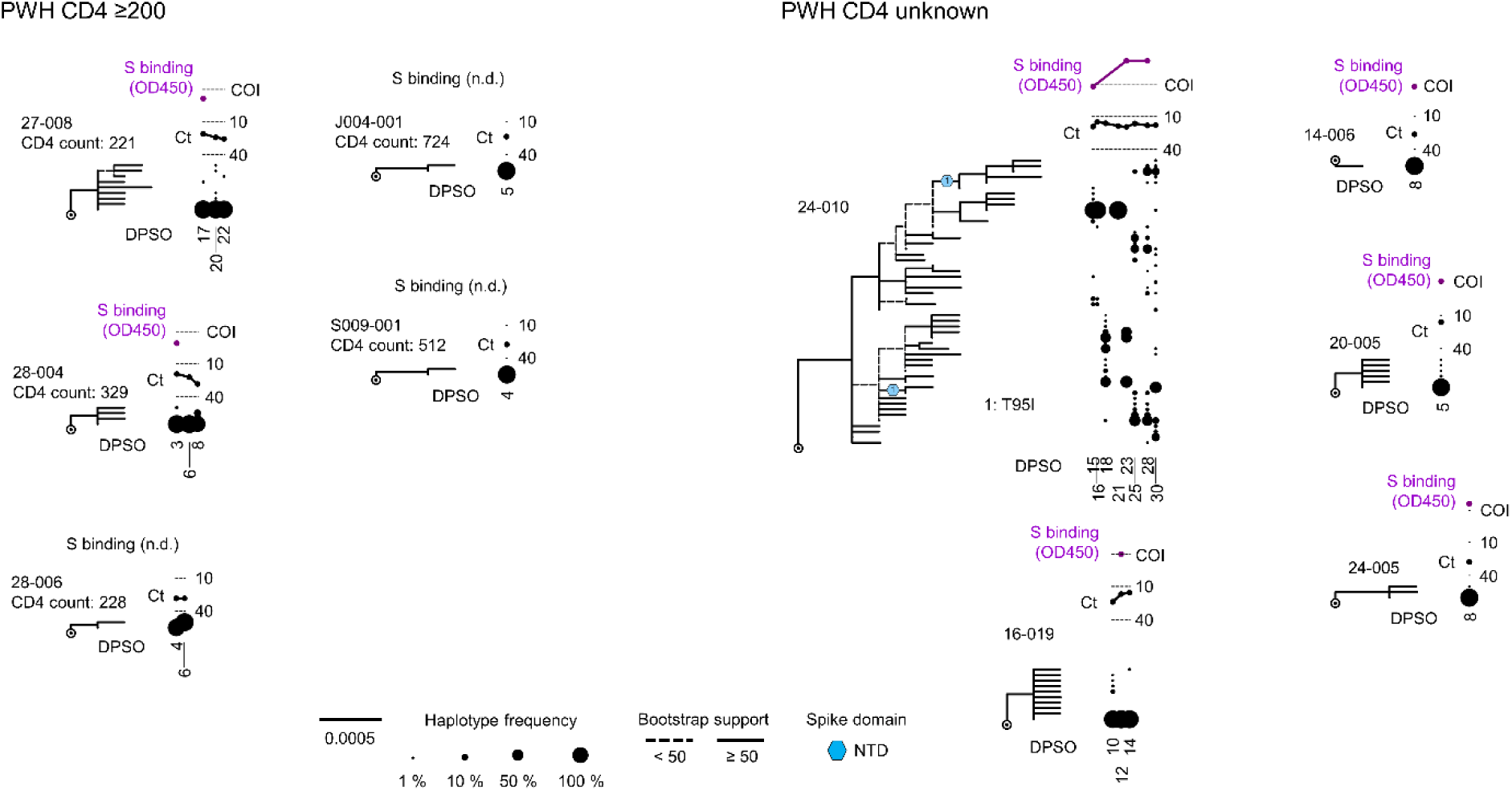
SARS-CoV-2 spike evolution in PWH with CD4 counts ≥200 cells/μL or unknown CD4 counts. Maximum-likelihood phylogenetic trees rooted on Hu-1 for all haplotypes from each PWH with CD4 count ≥200 cells/μL (left) or unknown CD4 count (right). Clades with bootstrap support less than 50% are indicated with dashed lines. Sites detected under positive selection within each participant (see Methods) are shown at their inferred location on the tree with numbered symbols; mutations corresponding to each number are listed beside each participant’s tree. Symbol shapes are coded by spike protein domain (see legend, center bottom). The frequency of each haplotype detected at each sample timepoint (in DPSO) in each participant is indicated to the right of the tree with a scaled black dot. SARS-CoV-2 RNA levels (rRT-PCR Ct values; black traces) and serum antibody binding to spike protein (optical density, 450 nm [OD450]; purple traces; n.d.-no data) are shown above the dot plot for each participant. The positivity cutoff index (COI) of 0.4 for serum antibody binding to spike protein is indicated with a dashed line.

**Extended Data Fig. 9.**
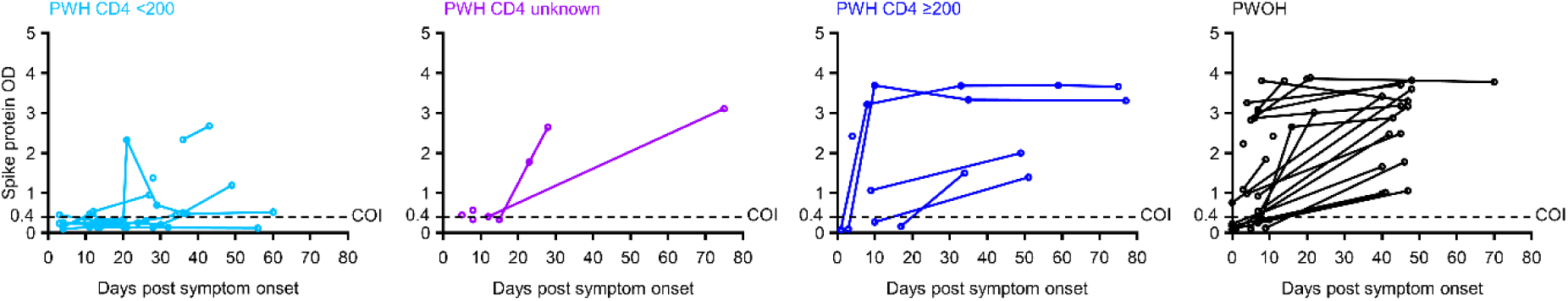
SARS-CoV-2 spike serum antibody binding responses in PWH and PWOH. Spike antibody binding titer over time in PWH subgroups and PWOH. The positivity cutoff index (COI) of 0.4 for serum antibody binding to spike protein is indicated with a dashed line.

**Extended Data Table 1.** Functional roles of mutations under positive selection in PWH with CD4 counts <200 cells/μL. Mutations detected as under positive selection (determined as in Methods) in at least one participant are shown. Presumptive functional roles were assessed by literature review as of September 2023. Immune evasion was defined as evidence of escape from any monoclonal antibody or from polyclonal sera; infectivity was defined as increased receptor binding, increased fusogenicity, or other growth advantage *in vitro*. Recurrently deleted regions RDR2 and RDR4 were included in the analysis despite being ineligible for dN/dS calculation via FUBAR because they were repeatedly observed in PWH with CD4 counts <200 cells/μL (see Fig. 3b) and were associated with frequency increases >20%.

| Variant | FUBAR<br>p > 0.90 | Frequency<br>Change | Infectivity | Immune Evasion | Additional notes |
| --- | --- | --- | --- | --- | --- |
| F4L | TRUE | TRUE | Unknown | Unknown | No references |
| L18F | TRUE | TRUE | No | Yes<sup>49</sup> |  |
| T19I/S | TRUE | TRUE | No<sup>50</sup> | Unknown | Decreased infectivity, unknown impact on neutralization |
| T22A | TRUE | TRUE | Unknown | Unknown | No references |
| T95I | TRUE | TRUE | Yes<sup>51</sup> | No | Potentially enables binding to AXL |
| D138H/N | TRUE | TRUE | Unknown | Unknown | Likely directional selection based on surveillance sequences<sup>52</sup> |
| RDR2 | N/A* | TRUE | No | Yes<sup>24,53</sup> |  |
| G142D/S | TRUE | TRUE | Unknown | Yes<sup>54,55</sup> |  |
| Y145D/H | TRUE | TRUE | Unknown | Yes<sup>56</sup> |  |
| RDR4 | N/A* | TRUE | No | Yes<sup>24</sup> |  |
| S247A | TRUE | TRUE | Unknown | Unknown | No references |
| S255P | TRUE | TRUE | Unknown | Yes<sup>57</sup> | S255F associated with loss of neutralization |
| G257D/S | TRUE | TRUE | Unknown | Unknown<sup>58,59</sup> | In antigenic supersite; effect on neutralization not significant for some tested NTD mAbs |
| N370K | TRUE | TRUE | Yes<sup>60</sup> | No |  |
| N440K | TRUE | TRUE | Yes<sup>61</sup> | Yes<sup>62</sup> |  |
| L452R | TRUE | TRUE | Yes<sup>63</sup> | Yes<sup>53</sup> |  |
| S477N | TRUE | TRUE | Yes<sup>64</sup> | Yes<sup>53</sup> |  |
| E484K | TRUE | TRUE | No | Yes<sup>53,65</sup> |  |
| S494P/T | TRUE | TRUE | Yes<sup>53</sup> | Yes<sup>53</sup> |  |
| N501Y/T | TRUE | TRUE | Yes<sup>53,66</sup> | Yes<sup>53</sup> |  |
| H655Y | TRUE | TRUE | Yes<sup>67</sup> | No<sup>68</sup> | Improved cleavage and fusogenicity |
| Q675R/L | TRUE | TRUE | Yes<sup>69</sup> | No |  |
| P681R/H | TRUE | TRUE | Yes<sup>70,71</sup> | Yes<sup>71</sup> | Escapes IFITM Restriction |
| A706V | TRUE | TRUE | Unknown | Unknown | No references |
| T716I | TRUE | TRUE | No<sup>72</sup> | No | Destabilization effect |
| N764K | TRUE | TRUE | Yes<sup>50</sup> | Yes<sup>50</sup> |  |
| G769V | TRUE | TRUE | Unknown | Unknown | Sporadic appearance, sometimes with E484K<sup>73</sup> |
| D950N | TRUE | TRUE | Yes<sup>74</sup> | No |  |
| A1078V | TRUE | TRUE | Unknown | Unknown | Also observed in<sup>75</sup> |
| E1092K | TRUE | TRUE | Unknown<sup>76</sup> | No | Improved stability of trimer |
| P1069S | TRUE | TRUE | Unknown | No |  |
| G1219V | TRUE | TRUE | Unknown | Unknown | Sporadic appearance globally<sup>77</sup> |

Pr(PS): Probability of positive selection
\*Indels not applicable to analysis with FUBAR

## Supplementary Table

**Supplementary Table 1.** Characteristics of study participants.

| HIV status | Study ID | Symptom onset date | Symptom onset ~ last pos (days) | First pos ~ last pos (days) | Number of samples sequenced | Age | Sex | Initial SARS-CoV-2 RNA (RT-PCR Ct) | CD4 count (cells/mL) | Plasma HIV RNA (copies/mL) | Cohort |
| --- | --- | --- | --- | --- | --- | --- | --- | --- | --- | --- | --- |
| PWH | 15-030 | 9/1/2020 | 79 | 76 | 12 | 33 | Female | 22.86 | 1 | 754,000 | Hospitalized |
|  | 16-036 | 10/4/2020 | 79 | 76 | 17 | 45 | Male | 21.47 | 1 | Unknown | Hospitalized |
|  | 15-029 | 8/27/2020 | 41 | 37 | 7 | 43 | Male | 20.32 | 6 | 4,290,000 | Hospitalized |
|  | 20-015 | 9/3/2020 | 20 | 16 | 8 | 37 | Female | 25.85 | 6 | Unknown | Hospitalized |
|  | 20-017 | 9/28/2020 | 37 | 33 | 10 | 52 | Male | 19.21 | 9 | 314,372 | Hospitalized |
|  | 30-014 | 8/6/2020 | 69 | 55 | 17 | 30 | Female | 24.65 | 18 | Unknown | Hospitalized |
|  | 16-033 | 8/21/2020 | 27 | 15 | 5 | 44 | Female | 20.00 | 25 | 167,000 | Hospitalized |
|  | 16-032 | 7/12/2020 | 58 | 22 | 2 | 36 | Female | 25.24 | 33 | 885 | Hospitalized |
|  | S006-001 | 11/22/2020 | 38 | 34 | 8 | 41 | Female | 32.56 | 43 | 3174 | Outpatient |
|  | 30-020 | 10/17/2020 | 20 | 10 | 5 | 41 | Male | 19.78 | 56 | 358,119 | Hospitalized |
|  | 16-037 | 10/5/2020 | 50 | 33 | 5 | 35 | Female | 19.80 | 71 | Unknown | Hospitalized |
|  | S074-001 | 7/3/2021 | 8 | 3 | 2 | 36 | Female | 30.09 | 136 | 151,722 | Outpatient |
|  | 20-005 | 7/5/2020 | 5 | 0 | 1 | 45 | Male | 18.87 | Unknown | Unknown | Hospitalized |
|  | 24-005 | 7/1/2020 | 8 | 0 | 1 | 39 | Male | 29.41 | Unknown | Unknown | Hospitalized |
|  | 14-006 | 8/10/2020 | 10 | 2 | 1 | 35 | Female | 28.40 | Unknown | Unknown | Hospitalized |
|  | 16-019 | 6/12/2020 | 19 | 9 | 3 | 51 | Female | 25.98 | Unknown | Unknown | Hospitalized |
|  | 24-010 | 8/24/2020 | 44 | 29 | 8 | 41 | Female | 22.40 | Unknown | Unknown | Hospitalized |
|  | 27-008 | 9/1/2020 | 38 | 21 | 3 | 30 | Female | 22.74 | 221 | Unknown | Hospitalized |
|  | 28-006 | 6/1/2020 | 8 | 7 | 2 | 58 | Female | 29.71 | 228 | < 400 | Hospitalized |
|  | 28-004 | 6/9/2020 | 10 | 7 | 3 | 51 | Female | 20.91 | 329 | Unknown | Hospitalized |
|  | S009-001 | 11/29/2020 | 7 | 3 | 1 | 38 | Male | 30.57 | 512 | 143 | Outpatient |
|  | J004-001 | 10/8/2020 | 10 | 5 | 1 | 47 | Female | 26.76 | 724 | < 400 | Outpatient |
| PWOH | J032-001 | 1/25/2021 | 5 | 3 | 1 | 42 | Female | 31.02 | - | - | Outpatient |
|  | J054-001 | 4/7/2021 | 7 | 5 | 1 | 47 | Male | 31.07 | - | - | Outpatient |
|  | S082-001 | 8/16/2021 | 7 | 5 | 1 | 66 | Female | 27.88 | - | - | Outpatient |
|  | 22-010 | 7/22/2020 | 9 | 2 | 1 | 45 | Male | 28.72 | - | - | Hospitalized |
|  | 30-013 | 8/2/2020 | 9 | 6 | 1 | 55 | Female | 28.18 | - | - | Hospitalized |
|  | 16-024 | 6/13/2020 | 11 | 0 | 1 | 49 | Male | 29.45 | - | - | Hospitalized |
|  | J041-001 | 3/7/2021 | 14 | 11 | 4 | 21 | Female | 22.35 | - | - | Outpatient |
|  | 16-016 | 6/9/2020 | 15 | 5 | 2 | 63 | Male | 25.26 | - | - | Hospitalized |
|  | 15-014 | 6/1/2020 | 18 | 10 | 3 | 30 | Female | 25.51 | - | - | Hospitalized |
|  | S013-001 | 12/11/2020 | 26 | 23 | 1 | 62 | Female | 22.74 | - | - | Outpatient |
|  | S048-001 | 4/26/2021 | 28 | 26 | 3 | 65 | Male | 25.42 | - | - | Outpatient |
|  | J039-001 | 3/4/2021 | 35 | 31 | 2 | 47 | Female | 25.62 | - | - | Outpatient |
|  | J039-006 | 3/14/2021 | 35 | 38 | 4 | 17 | Female | 24.99 | - | - | Outpatient |
|  | J039-008 | 3/9/2021 | 36 | 34 | 6 | 0.5 | Female | 26.29 | - | - | Outpatient |
|  | J039-003 | 3/13/2021 | 36 | 38 | 3 | 24 | Female | 25.24 | - | - | Outpatient |
|  | S015-001 | 12/14/2020 | 37 | 34 | 2 | 40 | Female | 26.52 | - | - | Outpatient |
|  | 29-004 | 6/7/2020 | 38 | 33 | 4 | 78 | None | 21.90 | - | - | Hospitalized |
|  | S039-001 | 3/3/2021 | 41 | 36 | 3 | 54 | Male | 27.88 | - | - | Outpatient |
|  | S014-001 | 12/15/2020 | 43 | 41 | 2 | 44 | Male | 26.69 | - | - | Outpatient |
|  | J060-001 | 4/19/2021 | 44 | 41 | 3 | 48 | Male | 22.25 | - | - | Outpatient |
|  | S023-001 | 12/17/2020 | 47 | 43 | 1 | 51 | Male | 31.46 | - | - | Outpatient |
|  | 29-008 | 7/31/2020 | 83 | 78 | 5 | 49 | Female | 31.97 | - | - | Hospitalized |
|  | J071-004 | asymptomatic | 7<sup>#</sup> | 7 | 2 | 48 | Male | 25.16 | - | - | Outpatient |
|  | J039-007 | asymptomatic | 23<sup>#</sup> | 23 | 3 | 15 | Male | 27.22 | - | - | Outpatient |
|  | J039-005 | asymptomatic | 25<sup>#</sup> | 25 | 3 | 17 | Female | 24.63 | - | - | Outpatient |

<sup>#</sup> For asymptomatic participants, date of first positive test was used as a proxy for date of symptom onset.

